# Electrostatic facilitation of odorant capture in insects

**DOI:** 10.64898/2026.02.04.703773

**Authors:** Alexander N. Borg, Beth H. Harris, Liam J. O’Reilly, Fraser A. Woodburn, David M. Withall, Ryan A. Palmer, Daniel Robert, József Vuts

## Abstract

Olfaction is a sensory modality common to most organisms. In insects, the primary olfactory organ is the antenna, where sensilla house olfactory receptor neurons adapted to detect volatile organic compounds (VOCs). Whilst olfaction is well-understood at molecular and neural levels, questions remain as to how, biophysically, airborne VOCs reach sensilla. Transport through passive diffusion and active antennal motion is empirically supported but cannot entirely explain the remarkably rapid VOC sampling rates. We present evidence that the insect antennae exploits electrostatic forces that amplify VOC transfer from bulk air to sensilla. In effect, charged antennae capture more ambient VOCs than neutral ones, also evoking an enhanced electrophysiological (EAG) response to VOCs. Experimentally altering the charge of isolated antennae modulates EAG responses and olfactory sensitivity. Multiphysics modelling incorporating electrostatic and fluid dynamic mechanisms supports empirical evidence. Altogether, this work reveals the existence of a previously unknown and complementary biophysical mechanism supporting olfaction.

**Graphical abstract:** 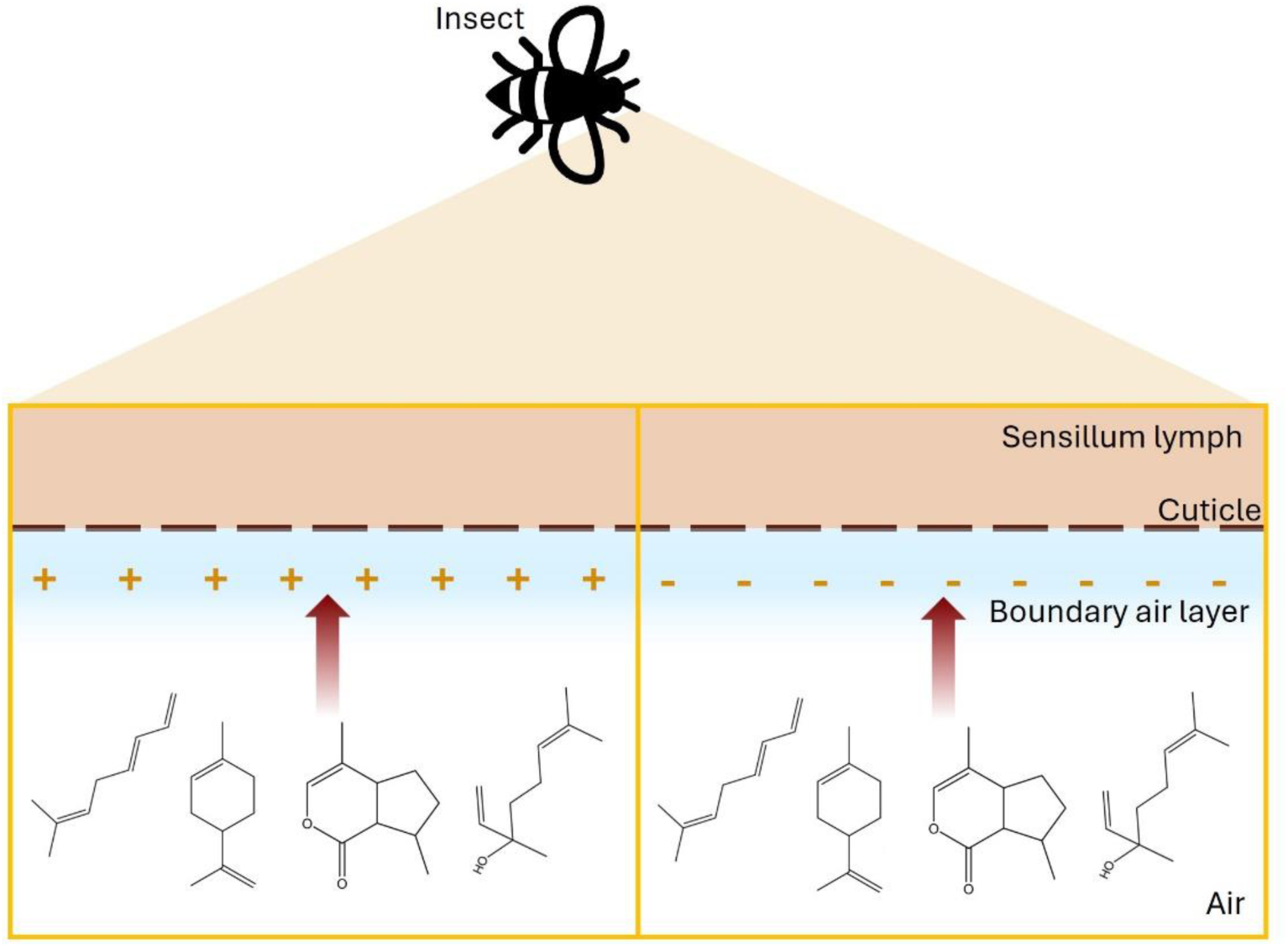

## Introduction

Olfaction underpins many important sensory ecological functions, from the localisation of prey, predators or mates to the deployment of pheromones to alert conspecifics and the identification of suitable oviposition sites. For terrestrial organisms, olfaction relies on the capture and detection of airborne scent molecules, i.e. volatile organic compounds (VOCs). Insects have the particularity of possessing externally facing olfactory systems, an arrangement substantially different from the internalised olfactory epithelia of most vertebrates. Here, olfactory sensory structures take the form of thousands of small porous sensilla, typically located on the insect’s antennae, inside which reside specialised olfactory receptor neurons (ORNs). The ORNs’ dendritic terminals present cellular membranes rich in odorant receptors (ORs), conferring sensitivity and specificity to VOC detection. The transfer of VOCs through the lymph surrounding dendritic terminals towards ORs is facilitated by odorant-binding proteins (OBPs), enhancing both reliability and speed of olfaction.^1^

Whilst olfaction is well-understood at the molecular and neural levels, questions remain regarding the coupling between dendritic terminals and VOCs residing in the aerial environment.^2^ Little is known on how fast and by what mechanism VOCs move through the boundary layer surrounding the insect antenna to reach, then enter, superficial pores on olfactory sensilla and subsequent ORs. To explain this very first step in the chain of olfactory events, current empirical and theoretical evidence suggest a combination between transport through passive diffusion and active antennal motion. Aimed at breaking and shedding the air boundary layer, antennal flicking motion enables a greater and faster availability of VOCs at the ORs. This mechanism receives support from empirical and theoretical work, whereby active antennal oscillations in the presence of VOCs serve to increase VOC capture rates in bumblebees.^3,4^ We propose here that this elegant process may not act alone, as it cannot entirely explain the extraordinary efficiency and rapid sampling rate observed in many insect species.^5–7^ In effect, recent fluid-dynamical evidence reveals that the convoluted antennal microscale morphology in moths enhances VOC capture and detection.^8^

Conventional olfactory theory states that reception of VOCs happens by chance via their diffusion towards the antennal surface.^9^ The finely branched comb-like antennae of male moths are estimated to adsorb about a third of molecules in an air stream,^10^ albeit such figure depends on many geometrical parameters not always known. In natural atmospheric conditions, this ratio is deemed to be much lower because of the patchy statistics of VOC concentration in heterogenous media.^11–13^ A salient limitation comes from the role played by the boundary layer surrounding antennal cuticular surfaces. It is estimated that the typical passive diffusive transit time through a thin 200 µm boundary layer is between 2-20 s for molecules with diffusion coefficients in the range of 1x10^-7^ – 1x10^-9^ m^2^/s,^3^ introducing both delayed responses and low dynamic range. However, the documented quick response time of the insect olfactory system to onset odorant stimuli, in the range of 3-10 ms,^14^ indicates a dynamic range exceeding that putatively dictated by passive buffering effects caused by the boundary layer.

We hypothesise that there is an additional and non-exclusive biophysical mechanism present, which enhances the sensitivity and temporal accuracy of olfaction in arthropods. Said mechanism would exploit the electrostatic forces arising between insects and their environment.^15^

Electric fields (E-fields) arise between electric charges and influence matter across a wide range of length scales: from subatomic particles, like electrons, through molecular and cellular structures to whole organisms, atmospheres, and even astrophysical environments. The electrostatic and electrodynamic interactions between charge-carrying particles largely dictate the chemistry of both the abiotic and biotic world, and thus consequently dictate the structure of life at many physical length scales. Of particular relevance here is that the distribution and mobility of charge within materials can influence biological and ecological processes.^15^ Indeed, recent work has highlighted the plethora of electrical interactions between organisms and their physical environment, demonstrating the complexity of their electric ecology.^16–19^

Arthropods accumulate surface charge on their cuticle as they move through their environment. Whilst the exact mechanisms of cuticle charging are unclear, triboelectrification is likely at work through friction between body parts such as wings, antennae, legs and hairs and any substrate or air. Hence, many animals living in terrestrial and aerial environments carry non-negligible electric charges.^15,20^ Often, but not exclusively, this charge is net-positive across the whole organism, resulting in an attractive force to sources of negative charge, owing to a Coulomb interaction.^21,22^ These charge differences facilitate ecological interactions, as observed with negatively charged pollen ‘jumping’ onto positively charged bees and butterflies prior to flower contact, enhancing pollination efficiency.^18,19^ Remarkably, it was shown in 1982 that the placode sensilla found on the antennae of honeybees hold a quasi-permanent electric charge. The author proposed that this charge attracts VOCs and thus enhances the capture efficiency of the olfactory receptor organ and improve its sensitivity,^23^ likely via dipole-dipole interactions. However, this putative mechanism has not been investigated further.

Considering the role electrostatics is poised to play in ecological relationships, we developed an alternative theory of olfaction that involves the electrostatic charging of sensory structures. Here, cuticular arthropod hairs endowed with charge interact with VOCs, themselves influenced by their dipole moment, a measure of the uneven distribution of charge held by a molecule. Notably, this interaction is predicted to occur outside the receptor at the sensillum and antennal level, influencing VOC capture, and is distinct from known nanoscale electrostatic attachment of volatile ligands to membrane-bound OBPs and ORs in the liquid phase.

Here, we propose that the electrically charged state of both the antennal cuticle and the olfactory sensilla increases VOC capture through attractive electrostatic forces (ex. Coulomb force) that overcome diffusion and advection timescales to aid the fast and efficient transfer of VOCs from air through the boundary layer to the sensory substrate.

## Results

We first measured the net charge on *Bombus terrestris, Aphidius ervi* and *Aphis fabae* antennae by dropping them into a Faraday cup. Measuring charge on *Drosophila melanogaster* antennae was not possible with our setup due to their extremely small size. Freshly amputated antennae, representing baseline measurements, showed a bias towards the negative range, but *A. fabae* antennae also bore positive values and thus had the broadest span of variation in charge (Figure 1). The amount of charge on *B. terrestris* antennae (Figure 1A) was estimated to be ca. two orders of magnitude higher than that of *A. ervi* and *A. fabae* antennae (Figures 1B and C). The placement of a neutralising plasma beam near the antenna generally decreased the otherwise large dispersion of charge values and reduced the amount of total charge for *B. terrestris* and *A. fabae* towards a more neutral state. Contact with the tungsten electrode also produced a narrower distribution of surface charge; here, application of 0 V caused clustering of values near zero in the positive range, a shift large enough to generate a statistical difference from the native (baseline) charged state of unbiased *B. terrestris* and *A. ervi* antennae. This effect became even more pronounced at the -8 V bias towards the positive range for all three species, which confirms that varying the electric potential influences the magnitude and polarity of antennal charge.

**Figure 1.**
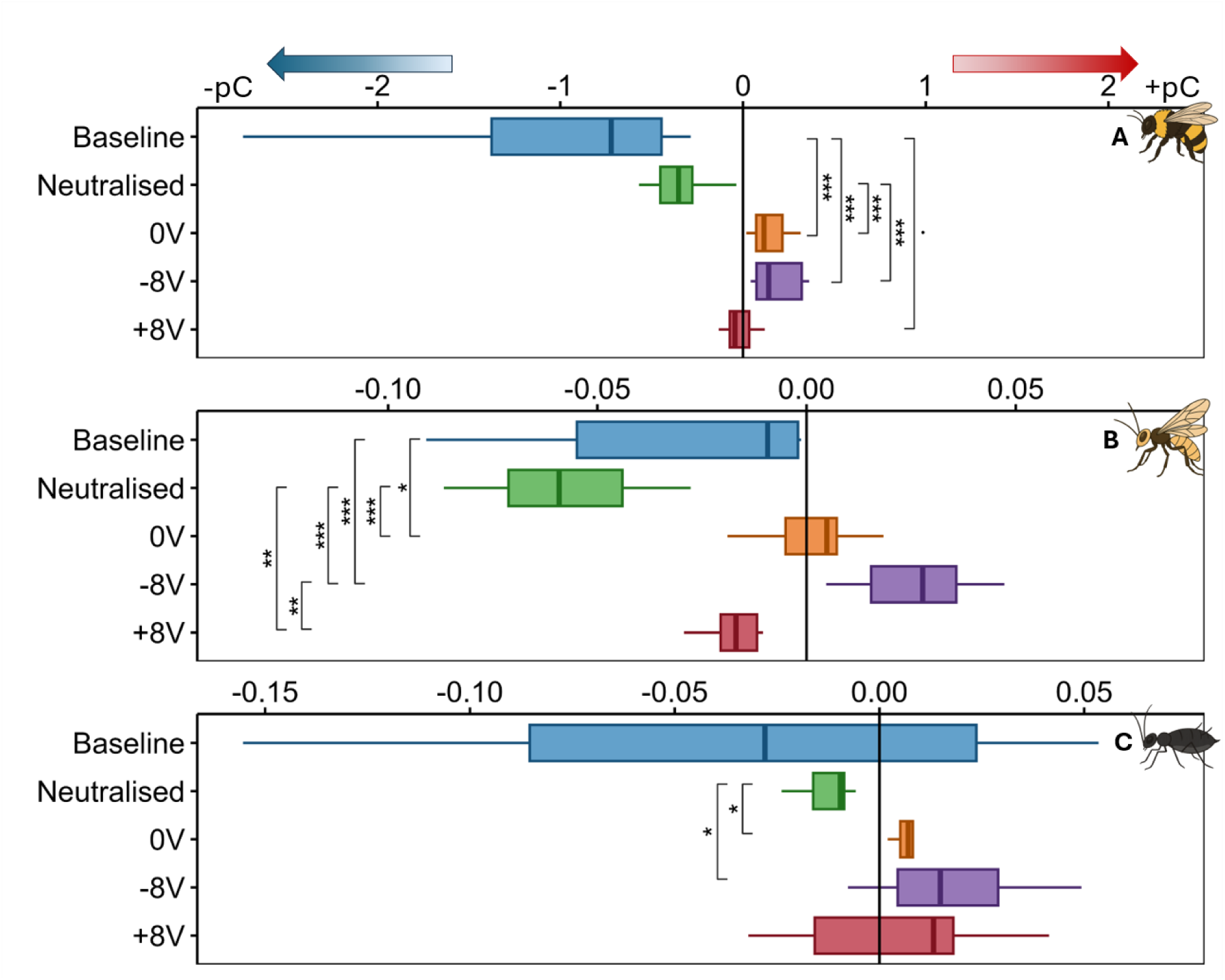
Net charge magnitude (pC) on *Bombus terrestris* (A), *Aphidius ervi* (B) and *Aphis fabae* (C) antennae, which were freshly amputated (Baseline), exposed to a plasma beam to reduce their spatial charge (Neutralised) or conductively treated with 0, -8 or +8 voltage (V) using a tungsten electrode. Measurement of net charge on individual antennae was made with a Faraday cup (n=10 antennae/species). Significance: · =P<0.1, *=P<0.05, **=P<0.01, ***=P<0.001, A and C: Kruskal-Wallis/Dunn test, B: ANOVA/Tukey.

A direct test of the role of electrostatics on olfaction was to assess the effect of surface charge on molecular adsorption from the surrounding air onto *A. ervi* antenna. By altering the surface charge of antennae using contact electrification, we found evidence of such effect for (4a*S,*7*S,*7a*R*)-nepetalactone and (*R*)-linalool, but not (*E*)-β-farnesene, where the -8 V bias increased the amount of adsorbed compounds as compared to 0 V bias (Figure 2). Under +8 V bias, (*E*)-β-farnesene showed a reduced, but non-significant, accumulation on the antenna from the airstream enriched with the compound.

**Figure 2.**
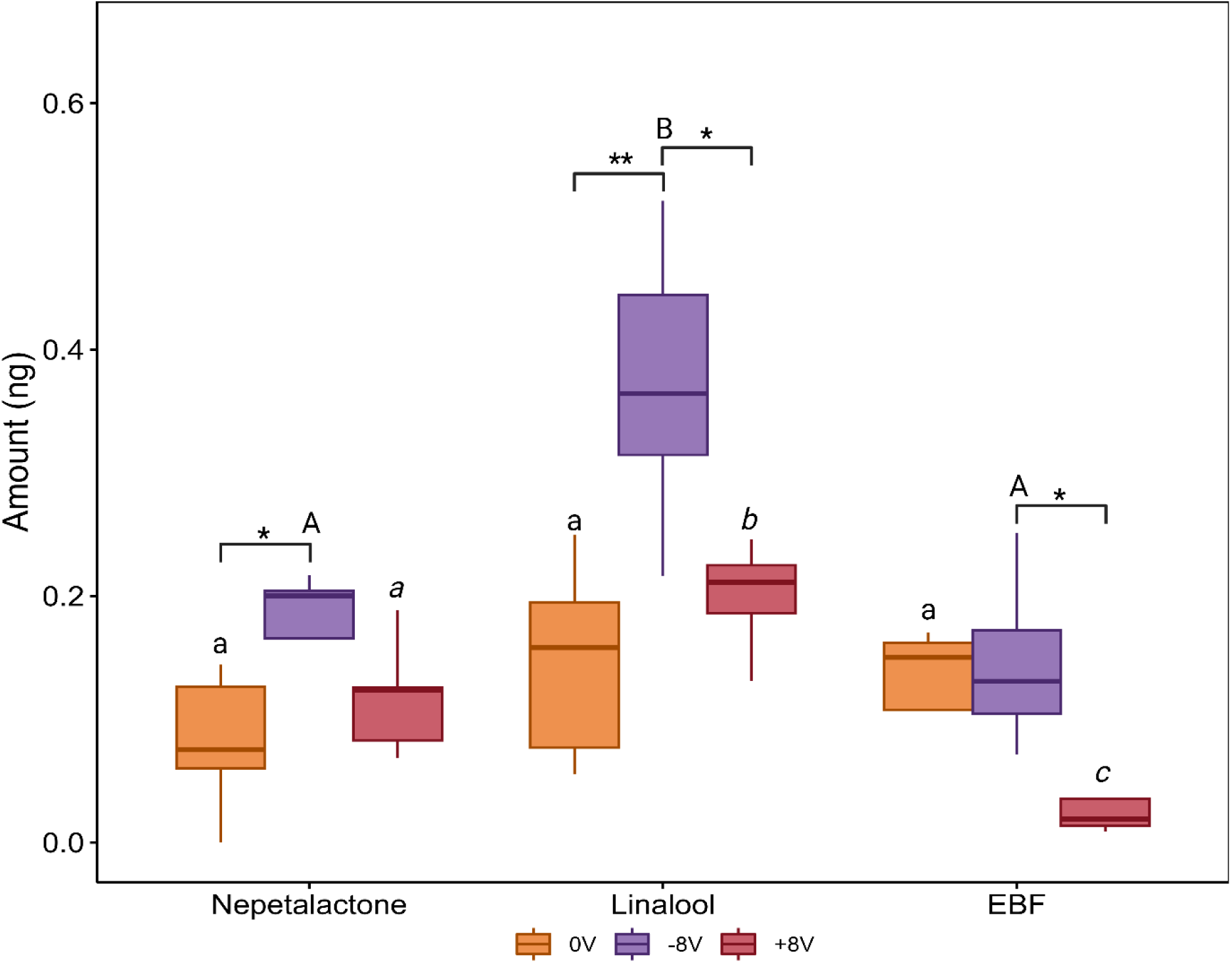
Amount of compound (ng) adsorbed onto *Aphidius ervi* antennae whilst exposed to 0, -8 and +8 V for 30 min (n=5 antennae/compound). Synthetic compounds (100 ug) were delivered to antennae through a constant stream of humidified air. Five antennae were extracted in diethyl ether after 30 min of exposure to make one replicate. EBF=(*E*)-β-farnesene. Significance within compounds: *=P<0.05, **=P<0.01 ANOVA/Tukey test per compound across electrical bias treatment. Significance within bias: lowercase, uppercase and italicized lettering, ANOVA/Tukey test per electrical bias treatment respectively across compounds.

Antennal electrophysiological (EAG) responses constitute clear evidence that test VOCs are detectable by the peripheral olfactory system. Consequently, we used the EAG technique to test the hypothesised link between antennal surface charge and VOC detection. Based on the observation that a plasma beam near the antenna reduces the overall antennal charge compared to its baseline state (Figure 1), we demonstrate that EAG responses to (*R*)-linalool become significantly smaller on *B. terrestris* antennae carrying reduced charge after plasma neutralisation than on antennae holding their baseline charge (Figure 3 A). The same effect was marginally significant on the *A. ervi* antenna (Figure 3 B).

**Figure 3.**
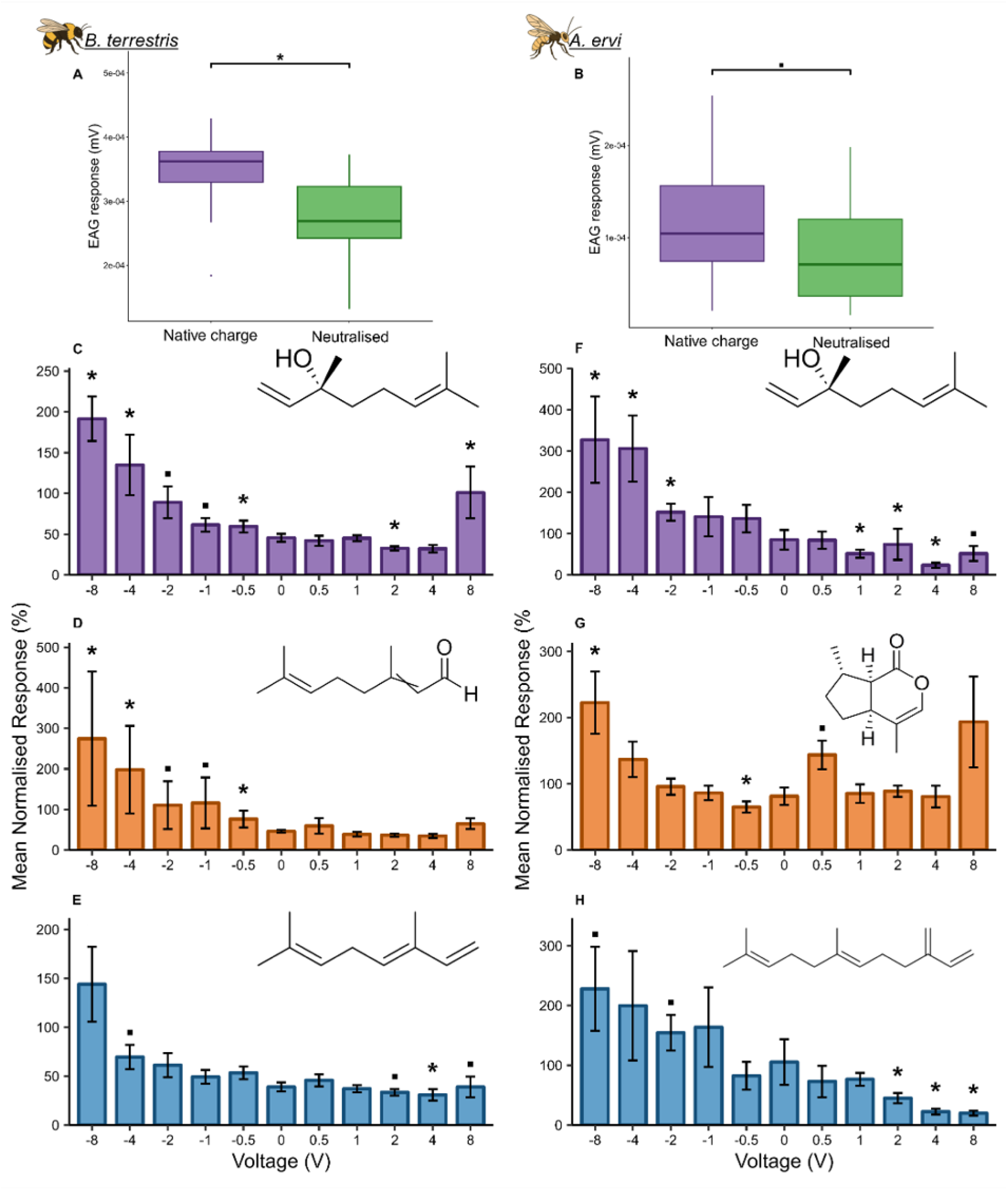
Effect of charge neutralisation via plasma beam on *Bombus terrestris* (A; n=11, dose=10 µg; Kruskal-Wallis/Wilcoxon test) and *Aphidius ervi* (B; n=24, dose=1 µg; Kruskal-Wallis/Dunn test) EAG responses to (*R*)*-*linalool. Electrophysiological (EAG) responses of *B. terrestris* and *A. ervi* antennae, exposed to a range of voltages, to synthetic compounds at a dose 10-fold lower than a significant EAG-active dose (baseline measurements Figure S1, mean ±SE). C: (*R*)-linalool (n=8, dose=1 µg), D: citral (n=7, dose=100 ng), E: (*E*)-ocimene (n=7, dose=10 ng), F: (*R*)*-*linalool (n=8, dose=100 ng), G: (4a*S,*7*S,*7a*R)-*nepetalactone (n=11, dose=100 ng), H: (*E*)-β-farnesene (n=8, dose=1 µg). EAG responses were normalised to a positive control: *B. terrestris* = benzaldehyde, *A. ervi* = (*E*)-caryophyllene. Charge was applied on antennae via tungsten electrode. Significance from diethyl ether solvent control: *·* =p<0.1, *=p<0.05, statistical tests used are described in Table S2. EAG responses from *Aphis fabae* and *Drosophila melanogaster* are shown in Figure S2.

Following manipulation of antennal charge with the plasma beam, we wanted to observe the effect of positive and negative electrical bias of the antennae on EAG responses. The EAG response increased across test compounds in a voltage-dependent and asymmetrical manner across a -8 — +8 V range, predominantly biased towards negative values (Figure 3C-H, Table S1, Figure S2). Furthermore, charge delivered with a -8 V bias, or occasionally weaker negative charges, increased antennal sensitivity (Figure 3C-H, Figure S2) when using ten-fold lower doses than the lowest EAG-active dose from baseline measurements (Figure S1). Table S1 show the same trend across the four species. Extending the range of positive electrical bias up to +12 V revealed a similarly increasing response pattern as observed in the negative range and highlighting the overall asymmetry of the effect of applied charge (Figure S3).

A consequence of facilitating availability of VOCs to the neural substrate would be to lower the sensitivity threshold to the concentration of volatile in the airflow. This was tested by presenting concentration series under a biased electrical regime. In effect, the -8 V bias lowered antennal detection thresholds across all model species and compounds down to doses on average four orders of magnitude below the lowest EAG-active dose on uncharged antennae (Figures 4 and S4, Table S1).

**Figure 4.**
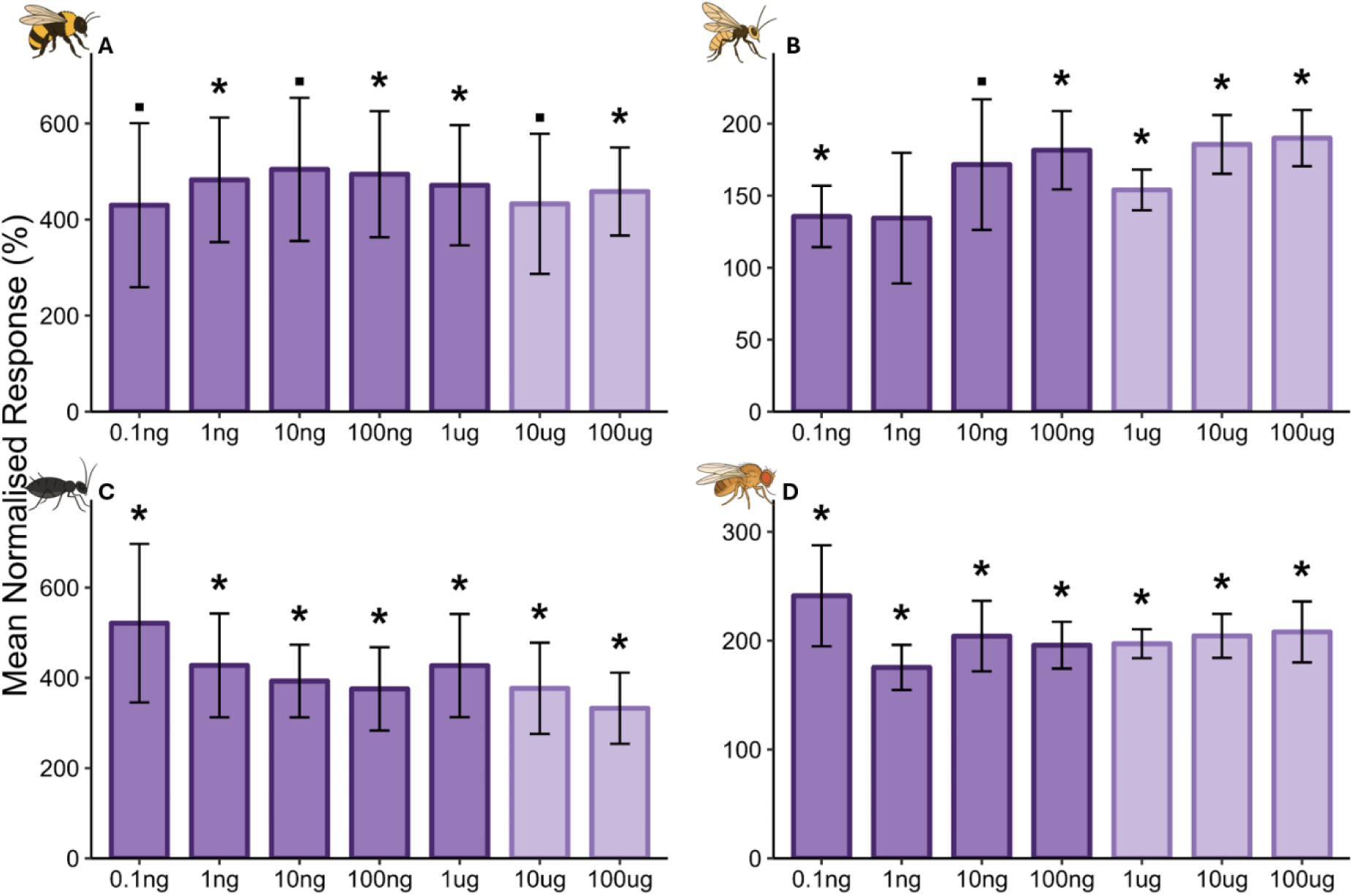
Electrophysiological (EAG) responses of *Bombus terrestris* (A, n=5), *Aphidius ervi* (B, n=5), *Aphis fabae* (C, n=7) and *Drosophila melanogaster* (D, n=5) antennae to a range of (*R*)*-*linalool doses whilst biassed with -8 V (mean ±SE). Significance from diethyl ether solvent control: *·* =p<0.1, *=p<0.05 (Student’s t-test, except for *B. terrestris* 100 µg: Wilcoxon test). Light purple bars represent the doses which induce a significant EAG response on uncharged antennae (baseline measurements, Figure S1).

To explore possible size and geometrical effects, we investigated the relationship between EAG response, antennal surface area and charge magnitude (as a function of voltage bias). There was a statistically significant interaction between EAG response, surface area and applied charge. EAG responses were higher on antennae with smaller surface area, in a descending order of *D. melanogaster, A. fabae, A. ervi* and *B. terrestris* (Figure 5A). This interaction was voltage-dependent, -8 V and +8 V biases showing the strongest influence. In correlation analysis between antennal charge density, applied charge and EAG response, a positive correlation between charge density and EAG response was observed (Figure 5B). Here, the -8 V treatment showed a significant difference compared to 0 V. Molecular dipole moment, on the other hand, showed no interaction with EAG responses under electrical bias, except for *D. melanogaster* (Figure S5).

**Figure 5.**
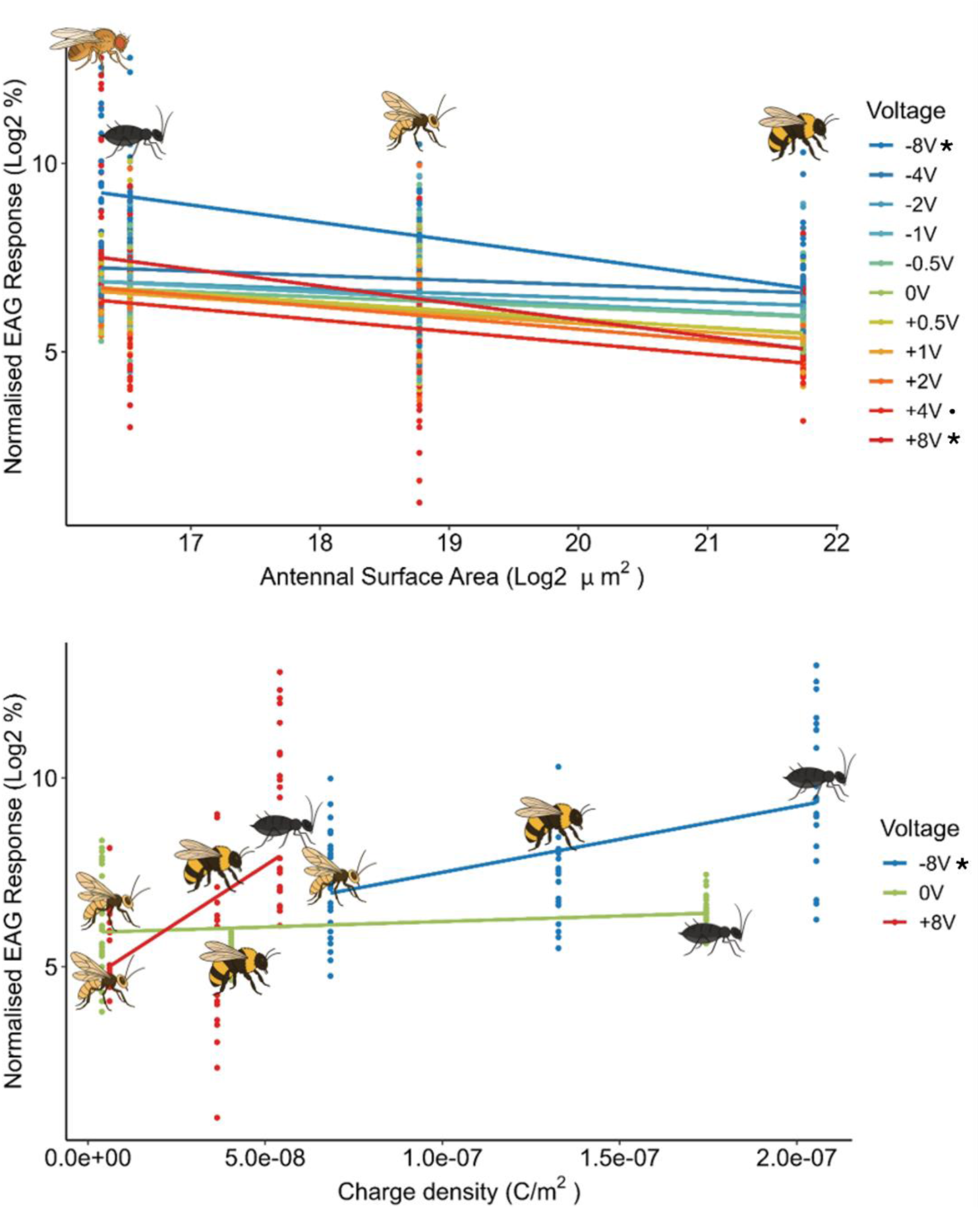
Relationship between electrophysiological (EAG) response and antennal surface area (µm^2^) (A), or EAG response and charge density (C/m^2^) (B) from antennae exposed to a range of voltages (V). Antennal surface area was calculated from stereomicroscopy images (n=3 antennae/species) and charge density calculated from charge measurement from figure 2. EAG responses were taken from *Bombus terrestris* exposed to 1 µg (*R*)-linalool, 100 ng citral and 10 ng (*E*)-ocimene (total n=21), *Aphidius ervi* exposed to 100 ng (*R*)-linalool, 100 ng (4a*S,*7*S,*7a*R)-*nepetalactone and 1 µg (*E*)-β-farnesene (total n=24), *Aphis fabae* exposed to 1 µg (*R*)-linalool, 100 ng (*E*)-2-heptenal and 100 ng (*E*)-β-farnesene (total n=24) and *Drosphila melanogaster* exposed to 100 ng (*R*)-linalool, 100 µg (*R*)-limonene and 1 µg (*E*)-2-hexenal (total n=22). Significance compared to normalised EAG response interaction at 0 V: *=P<0.05, *·* =P<0.1, GLM.

To investigate the possible electrostatic enhancement of the transport and capture of VOCs, a multi-physics mechanistic model was analysed. A dilute concentration of charged VOC particles is transported to the antenna via diffusion, convection and electrostatic migration. The considered charge values are several orders of magnitudes smaller than the fundamental charge to show the relative effect of antennal electrical fields, comparative to the order of magnitude effects polarised molecules experience in an electrical field.

For a hairless antenna in longitudinal flow, a boundary layer forms with the fluid velocity increasing from zero close to the antennal surface to the freestream value further away from it. For crossflow, a larger wake is seen past the antenna (Figure 6A and B, for 𝑈_∞_ = 0.1 𝑚𝑠^−1^). More biologically realistic models reveal that when hair sensilla are present, flow is further reduced near the surface, introducing mild fluid mixing between the hairs. Considering the capture rates given by (4) (Method details, Fluid-antenna interaction modelling) and only due to fluid dynamic effects, we find a higher capture rate over the hairless antenna in general (for longitudinal flow and crossflow) (Table 1). This partly results from local fluid flows replenishing the depleted concentration around the antenna and further confirmed by the general increase in capture with the oncoming flow rate. Overall, higher capture rates are seen in the longitudinal flow case. This is expected since the fluid flow, and thus concentration of VOCs, passes over a larger surface area and thus remain close to the antenna for a longer time, increasing the likelihood of capture.

**Figure 6.**
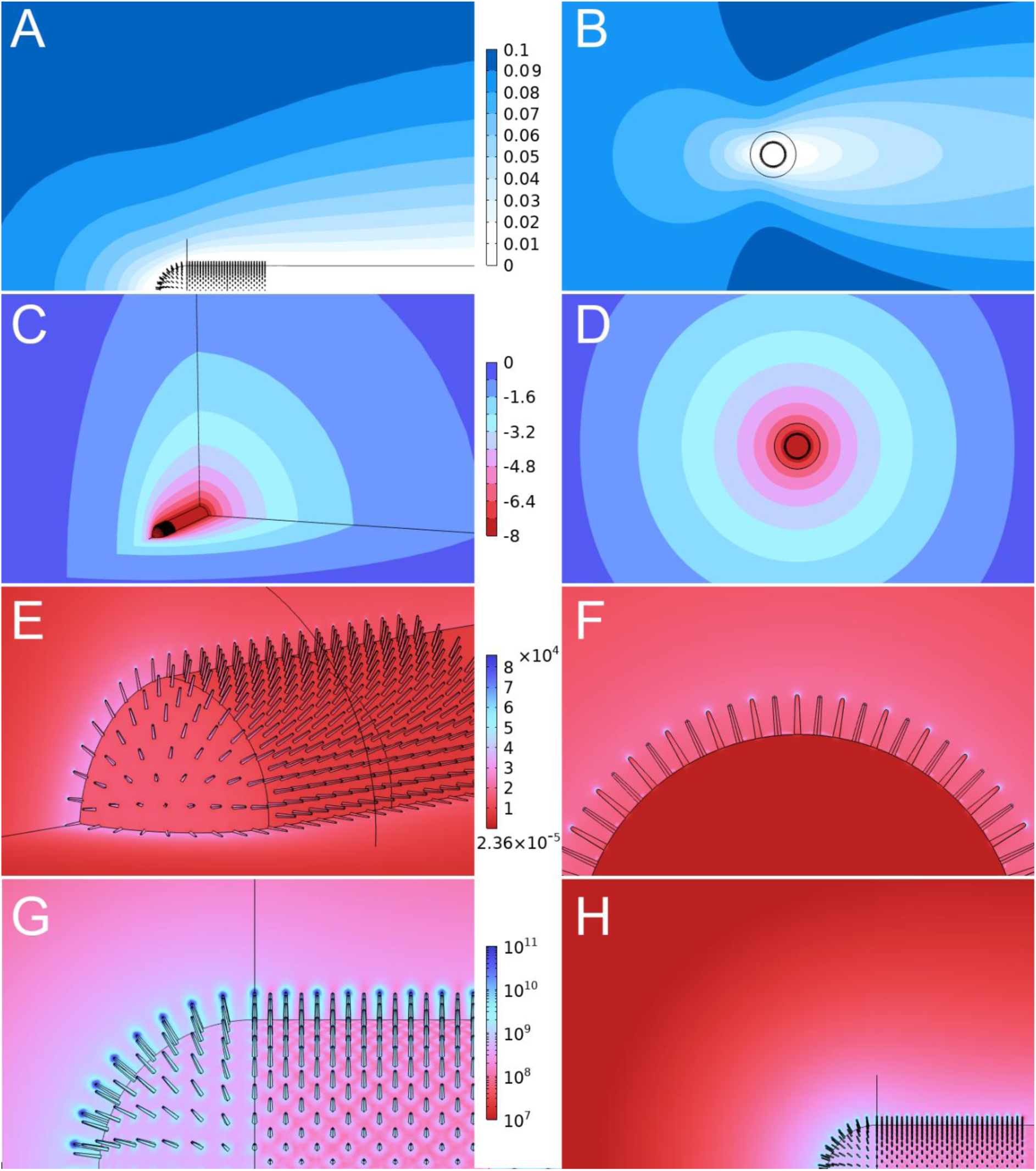
Fluid flow profile over an antenna for an incoming flow of 𝑈_0_ = 0.1 m/s. A boundary layer forms with slower flow speeds close to the antenna surface (tending to 0 at the surface) and increasing to the freestream velocity further from the antenna (A, B). Images of the modelled electric field generated by a biased antenna showing the electric potential decay from -8 V to 0 over several antennal lengths/radii (C, D) and the resulting electric field, showing enhancement at the hair tips and over the curved tip of the antenna, where two dimensions of curvature are present, up to O(1e5) in magnitude (E, F). The gradient of the electrical field around the antenna which acts on dipolar molecules, showing up to six orders of magnitude larger than the electrical field values (G, H).

**Table 1:** Capture rates, 𝐶_𝑎_, mol/s/m^2^, for molecules of zero charge and relative capture rates of non-zero charge molecules on an antenna with a dense hair coverage and no hairs for different flow speeds and morphology. There is a monotonic trend in capture rate with the effective charge of the VOCs, which is consistent across flow speeds. However, when the charge number is at least 0.1, the capture rate becomes invariant to flow speed, indicating that the electrostatic contribution to olfactory capture dominates transport forces due to fluid flow.

Longitudinal flow
| Charge Number | U = 0.001 m/s |  | U = 0.01 m/s |  | U = 0.1 m/s |  |
| --- | --- | --- | --- | --- | --- | --- |
| | Capture rates, $C_a$, mol/s/m<sup>2</sup> | | | | | |
|  | Dense | None | Dense | None | Dense | None |
| 0 | 0.0193 | 0.0276 | 0.0241 | 0.0344 | 0.0393 | 0.0558 |
| Capture relative to zero charge |  |  |  |  |  |  |
| 0.00001 | 1.0014 | 1.0015 | 1.0012 | 1.0012 | 1.0007 | 1.0008 |
| 0.0001 | 1.0145 | 1.0150 | 1.0117 | 1.0123 | 1.0073 | 1.0078 |
| 0.001 | 1.1505 | 1.1571 | 1.1209 | 1.1271 | 1.0745 | 1.0794 |
| 0.01 | 3.0185 | 3.1391 | 2.5502 | 2.6507 | 1.8810 | 1.9474 |
| 0.1 | 27.9813 | 29.6529 | 22.4226 | 23.7847 | 13.7939 | 14.6927 |

Crossflow
| Charge Number | U = 0.001 m/s |  | U = 0.01 m/s |  | U = 0.1 m/s |  |
| --- | --- | --- | --- | --- | --- | --- |
| | Capture rates, $C_a$, mol/s/m<sup>2</sup> | | | | | |
|  | Dense | None | Dense | None | Dense | None |
| 0 | 0.0068 | 0.0110 | 0.0111 | 0.0177 | 0.0217 | 0.0347 |
| Capture relative to zero charge |  |  |  |  |  |  |
| 0.00001 | 1.0019 | 1.0019 | 1.0014 | 1.0014 | 1.0006 | 1.0007 |
| 0.0001 | 1.0187 | 1.0190 | 1.0132 | 1.0135 | 1.0066 | 1.0068 |
| 0.001 | 1.1991 | 1.2033 | 1.1371 | 1.1403 | 1.0674 | 1.0691 |
| 0.01 | 4.1316 | 4.2002 | 2.8323 | 2.8816 | 1.8050 | 1.8279 |
| 0.1 | 42.1521 | 43.0508 | 26.2428 | 26.7818 | 13.4376 | 13.7185 |

**Table 2.** Mesh independence study showing relative error in capture rates for VOC capture with a finer mesh. Overall, all errors are less than 1%. The number of mesh elements depends on the number of hairs, hence the large number of boundary elements for the refine mesh in the longitudinal case.

| Relative error in capture rates | Boundary elements | $z_c = 0$ | $z_c = 10^{-5}$ | $z_c = 10^{-4}$ | $z_c = 10^{-3}$ | $z_c = 10^{-2}$ | $z_c = 10^{-1}$ | $z_c = 1$ |
| --- | --- | --- | --- | --- | --- | --- | --- | --- |
| Longitudinal | 26,215,931 | 0.3% | 0.3% | 0.29% | 0.25% | 0.05% | 0.65% | 0.67% |
| Crossflow | 5,809,087 | 0.17% | 0.17% | 0.18% | 0.2% | 0.4% | 0.8% | 0.67% |

Electrostatic forces and an effective charge on the VOC increase the overall capture rate of the antenna for both hairless and densely haired antenna and for all flow speeds (Table 1). The resulting electric field over the densely haired antenna is the same across test cases (Figure 6B-G). In Figures 6C and D, the electric potential varies from -8V on the surface to 0 in the far field. It must, however, be noted that the electric field is locally enhanced at the sharp hair tips and at antennal locations with high curvature following (2) (Figure 6E and F). In effect, high local electrical field strength associated with thin, sharp, high-curvature morphologies is an important element contributing to the presence, geometry and effectiveness of electrostatic forces.

The VOC capture rates are calculated in the presence or absence of charge on the antenna (Table 1). Remarkably, capture rates are independent from the presence of hairs. Also, we observe that effective charges as small as 0.01 q result in doubling or tripling or quadrupling of the overall capture rate, depending on flow speed. This result is in range of the experimental results for a -8 V biased antenna and represents the expected magnitude of electrostatic forces on polarised molecules close to the antenna (see below). Comparing to the influence of fluid flow, for an effective charge of at least 0.1 q, there is little variation in the capture results across flow speeds for both hairy and hairless geometries. Thus, the modelled regime shows the physical possibility for electrostatic forces to act independently, dominating fluidic transport and attracting more VOCs from the background dispersion to the antenna, at a faster rate than the fluid flow alone delivers. The action of electrostatic forces also results in an increased air volume from which VOCs can be captured, hence enhancing the sensing range.

Due to the small size of charges considered, the results are indicative of the order of magnitude effects expected for the action of an electrical field on polarized molecules. The force, 𝑭, experienced by an ionically charged molecule is proportional to the local electrical field, 𝑬, and the magnitude of the particle’s charge, 𝑞_𝑖_ = α𝑒, where 𝑒 is the fundamental charge and α a scaling constant, such that:

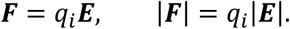

Here, 𝑭 is a vector associated with the direction and strength of the force on the particle. However, since a polarized molecule has no net charge, a force is generated by the gradient of the electrical field (i.e. how the electrical field varies in space) acting upon the molecule’s dipole moment, 𝒑 = 𝑞_𝑝_𝒅. Here, 𝑞_𝑝_ = 𝛽𝑒 with 𝛽 varying with the number of polarized charges in the molecule and 𝒅 = 10^−10^ 𝒅^ is a displacement vector defined by the spacing and alignment of opposite charges in the molecule, in the order of Angstroms ∼10^−10^ m. Thus, the force on a dipole in an electrical field is given by:

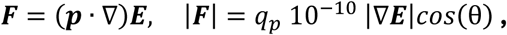

where θ is the angle of the displacement vector to the local electrical field gradient.

Considering the maximum force on a dipolar molecule (i.e. when the displacement vector aligns with the field gradient, 𝑐𝑜𝑠(θ) = 1), the magnitudes of the above forces are comparable if |∇𝑬|/|𝑬|∼(𝛼/𝛽)10^10^.

In Figures 6G and H, it is shown that the maximum value of |𝑬| is up to 1𝑒^5^, whilst the maximum of |∇𝑬| is several orders of magnitude larger at around 10^11^, hence |∇𝑬|/|𝑬|∼10^6^ at its largest. Thus, since we consider α = 1𝑒^−5^, 1𝑒^−4^, 1𝑒^−3^, 1𝑒^−2^, 1𝑒^−1^, our results cover the range of forces and interactions expected in the migration of polarized molecules in an electrical field for unknown 𝛽. Also, the field lines associated with gradient of the electrical field are similar in trajectory to those of the electrical field, towards the antennal surface. Finally, within one radius of the antennal surface (here, 0.1 mm away), the magnitude of the electrical field gradient remains up to 10000 larger than the electrical field magnitude, indicating the potential for equivalently large forces, as modelled here, on dipolar molecules within a region where we anticipate the electrostatic action to be most significant and effective.

## Discussion

In this study, we investigated whether electrostatics play a role in insect olfaction, thereby facilitating the capture of VOCs from the atmosphere through the boundary layer onto the sensillum cuticle. Across the species tested, our results reveal the variable ability of insect antennae to acquire and retain net electric charge. Application of charge onto the antennal surface increases adsorption of VOCs along with antennal EAG response. Altogether, our results demonstrate that electrostatics constitutes a functional element of olfaction, working at the interface between environment and sensor, enhancing sensitivity.

In the parasitoid wasp *A. ervi*, application of the -8 V bias increased the adsorption of both (4a*S*,7*S*,7a*R*)-nepetalactone and (*R*)-linalool, compared to the 0 V bias. This trend was also observed upon +8 V bias for (*R*)-linalool, however to a weaker and non-significant extent. This is opposed to (*E*)-β-farnesene, for which there was no significant difference in VOC adsorption across both a positive and negative 8 V bias, compared to 0 V, although at +8 V, a reduction was observed. It is known that VOCs with higher dipole moments are more strongly influenced by an electrostatically charged surface,^22^ offering an explanation for the increased adsorption of (4a*S*,7*S*,7a*R*)-nepetalactone and (*R*)-linalool on electrically biased *A. ervi* antennae, compared to (*E*)-β-farnesene, given their larger dipole moments. Also, the -8 V bias for (4a*S*,7*S*,7a*R*)-nepetalactone, and for (R)-linalool the +8 V bias, endows a higher magnitude of net charge than 0 V, which are likely to have a stronger polarising effect and hence induce stronger adsorption. Furthermore, the increased antennal accumulation of (4a*S*,7*S*,7a*R*)-nepetalactone and (*R*)-linalool at –8 V bias correlates with the increased EAG responses elicited for both compounds in *A. ervi*. The -8 V bias also somewhat increases the EAG response to (*E*)-β-farnesene, indicating that either even minor changes in its level of accumulation induce stronger EAG responses, or that other mechanisms are also involved. Under +8 V bias, the three compounds showed no significant change in accumulation compared to 0 V; (*E*)-β-farnesene levels, however, experienced a noticeable drop. These results corroborate EAG data, where a significant reduction in EAG response to (*E*)-β-farnesene is observed under the +8 V bias as compared to 0 V, but not for (4a*S*,7*S*,7a*R*)-nepetalactone and (*R*)-linalool, highlighting an electrophysiological asymmetry between negative and positive biases. Interestingly, (*R*)-linalool shows higher adsorption on *A. ervi* antennae than (4a*S*,7*S*,7a*R*)-nepetalactone at –8 V. Out of the three tested VOCs, (4a*S*,7*S*,7a*R*)-nepetalactone has the highest dipole moment and therefore, following our initial hypothesis and previous literature,^24^ it would be expected to be influenced to a greater extent by electric fields. This indicates the possible presence of a threshold from which the dipole moment of a compound is influenced by an electrically charged surface, increasing adsorption. Considering the humidified air stream within the experimental setup, the polar water molecules may antagonistically interfere with the test compounds in it, potentially influencing the electrostatic impact of the charged antennal surface. As (4a*S*,7*S*,7a*R*)-nepetalactone has the highest dipole moment of the three compounds, its interaction with water molecules is expected to be greater than that of (*R*)-linalool. Whilst these results highlight the role of electrostatic charge on antennal surface in attracting VOCs, they also point to different attraction between them, likely linked to their dipole moments interacting with local electric fields.

The propensity of insect antennae to exhibit surface charge is pivotal to the proposed mechanism of electrostatic attraction. Our results show that insect antennae readily hold electrostatic charge and that this charge can be manipulated using both contact electrification as well as through non-contact exposure to plasma. Testing functional relevance, we observe a general asymmetry in EAG responses towards negative electrical biases, with a higher positive bias (+10-12 V) required to induce a significant EAG response. This indicates that the insect antennal surface is more easily polarised positively in response to a negative electrical bias, which is consistent with the reduced VOC adsorption and lower antennal charge magnitudes recorded at +8 V bias compared to -8 V. Neutralisation *via* plasma beam decreases EAG responses; however, the difference in net charge elicited by electrostatic bias across the insect species highlights that the interaction between VOC, permanent/induced dipole moment and antennal surface charge is more complex than anticipated, with other factors, such as antennal morphology and local humidity, likely influencing interactions.

Charging behaviour varies between species, potentially reflecting intrinsic species-specific differences in electrical properties of their antennae. There are several potential mechanisms by which the electrode may generate or modify the antennal charge. These include: 1) dielectric polarisation, where internal and/or surface charges redistribute in line with an applied electric field; 2) tribolectrification through contact and/or friction, inducing electron transfer between cuticle and electrode; 3) adsorption of ions, particularly following surface potential changes; and 4) direct conductive charge transfer between the dielectric cuticle and/or haemolymph and its electrolytes and the electrode. While distinguishing between these mechanisms experimentally was beyond the scope of this study, the species-specific differences in charging behaviour provide insight into the electrical characteristics of insect antennae. It may be worth noting here that, as a general statement, it is phenomenologically very rare to find objects that do not present surface electric charges, a realisation of course also valid for all biological materials.

Both *B. terrestris* and *A. ervi* antennae were exclusively negatively charged, while *A. fabae* antennae presented a positive charge. Throughout all treatments, *B. terrestris* antennal charge was around an order of magnitude higher than the other two species, likely due to their larger size. These results suggest that variation in size and material properties of antennae between species may contribute to species-specific electrostatic properties, possibly underpinning different behavioural prerogatives.

The charging behaviour of antennae in response to plasma and voltage treatments further supports the notion of differing electrical properties between species. For instance, when exposed to plasma, the negative charge on both *B. terrestris* and *A. fabae* antennae was reduced compared to baseline measurements. Conversely, the magnitude of negative charge on *A. ervi* antennae increased on average. Exposure to plasma can neutralize the bulk charge of materials by producing large quantities of both positive and negative charge carriers, which adsorb to the material combining to result in a bulk charge approaching zero.^25^ This suggests that *A. ervi* antennae show a different and possibly polarity-specific ion affinity or adsorption behavior. Interestingly, contact electrification with 0 V brought the charge of all species closer to zero. In fact, the antennal charge polarity of all three species switched to positive on average, with only *A. ervi* antennae measuring negatively. This strongly suggest that there is a conductive pathway between the cuticle and the electrode, through which charge in the form of electrons is redistributed; however, other mechanisms such as surface interaction effects, like triboelectric charging, may also play roles in antennal charging behavior. Irrespective of the mechanism, it seems that there are clear species-specific charging behaviors in response to both contact and non-contact electrification, likely to be the result of inherent differences in antennal electric properties.

The most striking difference in antennal charging is seen in response to the ±8 VDC treatments. Both *B. terrestris* and *A. ervi* antennae mirror the polarity of the treatment potential. A positive electrode potential results in a negatively charged antenna and vice versa. Such a response suggests that the antennae of these species electrostatically polarize with respect to the potential of the electrode and its incident electric field. External and/or internal polar molecules or charge carriers orient or move in relation to the electric field produced by the electrode. Removing the source of the electric field will cause the displaced charges to reorient; however, in non-conductive dielectric materials this is not an instant process, resulting in the apparent charge of the material persisting in time.^26,27^ Following removal of the electrode, we measured the charge quickly enough to capture these polarization effects within a few seconds, suggesting the antennae possess distinct dielectric properties.

*Aphis fabae* antennae behaved differently, charging positively on average in response to both positive and negative DC potentials, likely indicating fundamentally different electric properties and thus modes of charging. It is apparent that *A. fabae* antennae are more conductive than those of *B. terrestris* and *A. ervi*, resulting in the antenna equalizing to the electrode’s potential upon contact. If the antenna were a perfect conductor, it would be expected to have a charge close to zero once the electrode is removed, as charge almost instantly redistributes.^27^ However, the measured residual positive charge, regardless of electrode polarity, suggests that the antenna loses electrons following contact with the electrode. Such results may hint at a combination of conductive and triboelectric charging. Furthermore, in *A. fabae*, *A. ervi* and *B. terrestris*, baseline antennal charge density did not directly scale with antennal surface area (Figure S6). Specifically, the average baseline antennal charge density of *B. terrestris* was not found to be lowest in magnitude, as would be predicted if charge density scaled with antennal surface area alone, suggesting that across these three species, antennal charge regimes are likely influenced by additional properties of the antenna and may reflect adaptive variation. It is important to note that studying the electrical properties of insect cuticle is challenging. The cuticle acts like a dielectric in some species, a semi-conductor in others and may seemingly show conductive properties,^28^ as observed in *A. fabae*. In effect, it can be suggested here that cuticle is a heterogenous polymer exhibiting dielectric characteristics such as polarization. The mechanisms of charging discussed here are speculative and non-exhaustive. This study did not set out to elucidate these phenomena; nevertheless, it would certainly be a useful and fruitful area of future research.

Insect antennae are morphologically very diverse,^29^ housing a range of different sensillum types with different densities and distributions. This is evident across the four species investigated,^30–32^ with marked differences arising even between species of the same order (Hymenoptera: *B. terrestris*, *A. ervi*).^30,31^ Such microanatomical differences likely influence the local electric field distribution of the antenna; for example, in male honeybees, antennal placode sensilla have been found to hold a different electrostatic charge from the surrounding cuticle.^33^ Other studies have also suggested that insect sensilla and cuticle carry electrostatic charges.^23,34^ Whilst, crucially, the local electrostatic properties of the antennal sensilla of the four species investigated herein are not known, it is possible to speculate that antennal surface morphological factors, such as sensillum density and aspect ratios, may influence the bulk charge of the antenna under natural conditions, upon charge neutralization or voltage bias application. Additionally, ultrastructural differences, such as cuticular thickness, may play a role in the capacity of the antenna to acquire, maintain or dissipate charge. Chitin is a major component of the exoskeleton of insects, present also in the antennal cuticle.^35,36^ The triboelectric chargeability of chitin and its varied forms are documented^37,38^ and for this reason, forms of chitin are utilized as a dielectric material for triboelectric nanogenerators, which convert mechanical energy into electrical energy via triboelectrification.^37–39^ Whilst in this study, antennal charges were manipulated by the application of DC voltage biases, variation in the electrical response of the antennae both between species and treatments (Figure 3) may be partially attributable to ultrastructural factors, such as the form and content of chitin present in the cuticle, its thickness and antennal aspect ratio. Since our results show that manipulation of the antennal charge state affects olfactory sensitivity, morphology and ultrastructure may in turn modulate the charge properties of the antenna. Exploring this interplay represents a valuable direction for research.

Insect cuticles also contain superficial cuticular hydrocarbons and proteins within the chitin matrix, whose composition are typically unique across genera and between species,^40,41^ with specific ecological functions such as contact sex pheromones in the case of cuticular hydrocarbons in parasitic waps.^42,43^ In fact, chemical analysis of insect cuticular hydrocarbons have shown that composition across bumble bee (*B. terrestris*),^44,45^ parasitic wasps within the Braconidae family,^42,43,46^ and aphids (*Aphis gossypii* Glover)^47^ differ between each other. This may act as another factor influencing the electrostatic characteristics of the cuticle across insects. Composition differences likely affect the polarizability of the cuticle due to the different functional groups present across surface hydrocarbons and proteins, creating dipole moments across the molecule and on a wider scale across the insect cuticle. This characteristic may be analogous to the ordering of inorganic materials across a triboelectric series.^48^

The modelling analysis revealed how antennal capture rates of minimally charged VOCs increase with antennal charge, representing the order of magnitude effects expected for dipolar molecules. Capture rates were predicted to at least double for lower charge numbers commensurate with the experimental results (e.g. charge numbers of 0.01 q). Additionally, higher VOC capture on the antenna occurred with higher flow speeds for both uncharged and charged VOCs. Increased capture from a faster fluid flow occurs due to more VOCs being brought closer to the antennal surface acting to replenish the depleted concentration around the antenna. Interestingly, at charge numbers higher than 0.01 q, the VOC capture rate did not vary between low and high flow speeds. Similarly, when the flow speed is at its lowest (0.001 m/s), the antenna has a mildly higher VOC capture rate. In each case, it is deemed here to be due to electrostatics augmenting the delivery of VOCs by the fluid flow, increasing capture. This computational result thus established the physical possibility of an electrostatic enhancement of olfactory capture, whereby VOCs are charged significantly to a threshold at which the electrostatic forces dominate the capture process. Here, in effect, the electrostatics attract VOCs faster than the fluid delivers, showing no variation in capture rate with flow speed. Notably, these values are closer to those of ionic charging than those associated with the molecular dipole moment regime. In theory, the magnitude of electrostatic force acting on a polarised molecule will increase with its proximity to the gradient of the electrical field increasing by several order of magnitude in comparison to the electric field itself.

For a biased antenna modelled at a fixed surface potential of -8 V, the electric field, and thus its gradient, on and local to the antenna is shown to concentrate on the sensilla and at the curvature of the antennal tip, with the electric field gradually dissipating outwards across several antennal lengths. In studying both dense arrays of sensilla and no sensilla, we found that the overall VOC capture rate was marginally higher in the hairless case. This, in part, is shown to be due to the hairs slowing the fluid flow around the antenna, thereby reducing the local replenishment of VOCs. Our data collectively suggest that electrostatics can significantly enhance olfactory capture for polarisable molecules and even dominate potential fluid flow forces for ionically charged particles by attracting more from the background dispersion to the antenna and at a faster rate.

In effect, electrostatic bias on an antenna increases VOC capture and induces stronger EAG responses. Our empirical and theoretical analyses together provide evidence that biologically relevant electrostatic forces have the capacity to determine VOC capture rates by antennae in conjunction with fluid flow. This electrostatic effect can also be partially influenced by the dipole moment of the VOC. We show that the proposed electrostatic mechanism is likely to be common across insect genera; however, its prevalence may be dependent on the sensory biology and ecology of the insect, opening up enticing avenues in the biophysical studies of olfaction and its diversity.

### Limitations of the study

To better understand olfaction in natural settings, the effect of electrostatic forces in turbulent air flows need to be characterised, which was not part of this study. VOCs carried by turbulence are poised to encounter the antenna in packets with their own spatial and temporal statistics, ^11,49,50^ whereby electrostatic forces compete with air turbulence to capture various volatile semiochemicals, enabling appropriate behavioural decisions. Furthermore, measuring the antennal charge magnitude of *D. melanogaster* antennae was not possible due to their small size, which will require a modified version of the Faraday cup. In our model, we can only prescribe a fixed charge on the VOCs. If their effective charge is the result of polarisation and hence a dipole moment, their polarity and effective charge magnitude can be expected to vary during their trajectory. In this instance, we anticipate that VOCs would take the opposite sign to the background field and hence only attractive forces will be at work. The precise physics of such a dynamic process require further experimental validation to qualify this hypothesis. In real terms, effective charge is likely to be context-dependent, resulting in the interplay between atmospheric ionic composition and varied dielectrics moving at speed through it. Presently, the complexity of electrostatic phenomena needs highlighting, along with their increasingly apparent roles in the life of arthropods, other animals, plants and possibly all life. Thus, a further point of discussion is the assumption of independence between the fluid regime and electrical field generation. Whilst valid for disperse and diffuse VOCs, the possible role of ions in the charging of an antenna *in vivo* and the possible triboelectric charging of an antenna in an air flow require investigation. Both mechanisms may serve to enhance or weaken the strength of its electrical field within different modes of transfer.

### Resource availability

#### Lead contact

Requests for further information or access to resources not already publicly available should be directed to the lead contact, Dr. József Vuts and Daniel Robert -

## Supporting information

supplemental text - specimen preparation and fixation methods for bioimaging

## Materials availability

No unique materials or reagents were generated in this study Data availability

All generated data are available from the Rothamsted Research repository (DOI: https://doi.org/10.23637/ocxemf97)

## Acknowledgments

We would like to thank Kirsty Halsey and the Rothamsted Research Bioimaging team for carrying out the sectioning and imaging of insect antennae in this study. This work was funded through a BBSRC Pioneer award (BB/Y512886/1) to J.V. and D.R. F.A.W was funded by a BBSCRC SWBio studentship grant. J.V. and A.N.B. acknowledge support from the Rothamsted Research Growing Health Institute Strategic Programme [BB/X010953/1; BBS/E/RH/230003A]. B.H.H., L.J.O, R.A.P and D.R. were supported by an advanced grant from the European Research Council (ERC-ELECTROBEE 743093) to D.R.

## Author contributions

J.V. and D.R. were responsible for funding acquisition and conceptualisation. J.V., D.R., L.J.O., B.H.H. and F.A.W were involved in acquisition of preliminary data and initial method development. J.V., D.R., L.J.O., B.H.H. and A.N.B participated in experimental design and method development. J.V., L.J.O. and B.H.H. supported A.N.B. in the execution of some experiments. A.N.B. carried out data analysis and visualisation. R.A.P developed and carried out multiphysics and molecular modelling and analysis. D.M.W synthesized (*E*)-β-farnesene and (4a*S*,7*S*,7a*R*)-nepetalactone. J.V., D.R., L.J.O., B.H.H., A.N.B. and R.A.P drafted the manuscript. All authors drafted the manuscript, reviewed it, and agreed on its contents towards submission.

## STAR Methods

### Key resource table

| REAGENT or RESOURCE | SOURCE | IDENTIFIER |
| --- | --- | --- |
| **Chemicals, peptides, and recombinant proteins** |  |  |
| Diethyl ether | Fisher Scientific | 10368182 |
| (4a*S*,7*S*,7a*R*)-Nepetalactone | Synthesized in-house | CAS: 21651-62-7 |
| (*E*)-β-farnesene | Synthesized in-house | CAS: 18794-84-8 |
| (*R*)-linalool | Botanix Ltd. | CAS: 126-91-0 |
| (*E*)-2-heptenal | Fluka Ltd. | CAS: 18829-55-5 |
| (*E*)-β-caryophyllene | SAFC | CAS: 87-44-5 |
| (*E*)-ocimene | Synthesized in-house | CAS: 3779-61-1 |
| citral | Fluka Ltd. | CAS: 5392-40-5 |
| benzaldehyde | Sigma-Aldrich | CAS: 100-52-7 |
| (*R*)-limonene | Fluka Ltd. | CAS: 5989-27-5 |
| (*E*)-2-hexenal | SAFC | CAS: 6728-26-3 |
| (+)-fenchone | Fluka Ltd. | CAS: 4695-62-9 |
| **Experimental models: Organisms/strains** |  |  |
| *Aphis fabae* | Rothamsted Farm (co-ordinates: 51°48'36.8"N 0°22'34.4"W) | N/A |
| *Aphidius ervi* | Koppert Uk Ltd. | 02720 |
| *Bombus terrestris* | Biobest Uk Ltd. | 006219 |
| *Drosophila melanogaster* | Blades Biological Ltd. | LZJ 398 |
| **Software and algorithms** |  |  |
| MATLAB R2022a | MathWorks Inc. | https://uk.mathworks.com/products/matlab.html |
| National Institute of Standards and Technology (NIST) mass spectral library 2020 | National Institute of Standards and Technology | https://chemdata.nist.gov/dokuwiki/doku.php?id=chemdata:st art#libraries |
| Syntech EAG software package EAG v1.0 (6/1993) | Syntech | N/A |
| COMSOL Multiphysics software V 6.2 | COMSOL inc. | https://www.comsol.com/comsol-multiphysics |
| RStudio V2023.12.1+402 | Posit PBC | https://posit.co/download/rstudio-desktop/ |
| R V4.3.3 | The R Foundation<sup>58</sup> | https://cran.r-project.org/bin/windows/base/old/ |
| **Other** |  |  |
| USB-6009 NI DAQ | National Instruments Corp. | 779026-01 |
| Plasma Lighter | VEHHE™ Luoyang Hengshu shangmao Co. Ltd. | B0DN61MSMF |
| FG100 Function generator | Walfront | B08FYM7D4R |
| Agilent 8890 series GC | Agilent Technologies | https://www.agilent.com/en/product/gas-chromatography/gc-systems/8890-gc-system?srltid=AfmBOorst-COCyUqa7stR1d7755u6zmjffElhFcndqSeUz2NxAbvQajy |
| HP1 column GC column | Agilent Technologies | 19091Z-115E |
| Agilent 5977B GC-MSD | Agilent Technologies | https://www.agilent.com/en/product/gas-chromatography-mass-spectrometry-gc-ms/gc-ms-instruments/5977c-gc-msd |
| Electroantennography (EAG) system | Syntech | N/A |
| Faraday Cup | (Harrison & Robert,<sup>51</sup>) | N/A |
| Zeis Axioimager Z2 with Zeis Axiocam 512 colour camera | Carl Zeiss AG | https://www.zeiss.com/microscopy/en/products/light-microscopes/widefield-microscopes/axio-imager-2-for-life-science-research.html |

### Experimental model and study participant details

*Aphis fabae* Scopoli, originating from Rothamsted farm (Hertfordshire, UK, co-ordinates: 51°48’36.8”N 0°22’34.4”W), were reared on *Vicia faba* L. cv. “The Sutton” in ventilated Perspex cages at 20°C, 60-70% humidity and 16:8h light:dark regime. *Aphidius ervi* Haliday were purchased from Koppert UK Ltd. (Suffolk, UK) and stored at 5°C until use. *Bombus terrestris* L. were purchased as hives from Biobest UK Ltd. (Kent, UK) and kept at 20°C. Wild-type *Drosophila melanogaster* Meigen were purchased from Blades Biological Ltd. (Kent, UK) and stored at 20°C.

### Method details

#### Faraday cup charge measurements

Antennal charge measurements across different treatments were recorded using a recently developed and described Faraday cup setup^51^. The Faraday cup was placed within a larger Faraday cage for electrical isolation and connected to a computer via a data acquisition module (NI USB-6009, National Instruments Corporation, Austin, TX) to retrieve charge readings via MATLAB R2022a (MathWorks Inc., Nattick, MA), scripts provided in supplementary (Supplementary text 1). Insect antennae were excised using a scalpel under a stereomicroscope (model M7A Wild Heerbrugg, Switzerland) and placed on a wooden stick, which was fastened to a micromanipulator using non-conductive adhesive putty and manoeuvred so that the antenna was directly over the Faraday cup opening. For the “Baseline” treatments, antennae were directly pushed into the Faraday cup using another wooden stick, recording the antennal charge. Wood was used as the material due to its electrically insulating properties and its near-neutral position on the triboelectric series, thus minimising its influence on antennal charge properties. For the “Neutralised” treatment, the antennae were treated with pulses from a plasma lighter (VEHHE^™^ DHQHS, Luoyang Hengshu shangmao Co. Ltd., China) by pressing the “on” button of the lighter at a 5 mm distance from the mounted antenna to provide a 2 s plasma discharge, eight times at 1 s intervals, prior to being dropped into the Faraday cup. A tungsten wire attached to an FG-100 DDS function generator (Walfront, China) and fixed on a micromanipulator was manoeuvred to touch the antennae and impart 0 V, +8 VDC and -8 VDC treatments, prior to being dropped into the Faraday cup. This was carried out for *A. fabae*, *A. ervi* and *B. terrestris* antennae, collecting ten replicates per treatment and species.

#### VOC antennal adsorption

Freshly excised *A. ervi* antennae were suspended at their bases on a glass electrode filled with Ringer solution (without glucose) and attached to a micromanipulator, as described in fig. S1. A tungsten electrode, fixed to a micromanipulator and connected to an FG-100 DDS function generator, was positioned to be touching the centre of the antenna from behind to not obstruct the flow of VOCs toward the antennae. Using the function generator, 0 V, +8 VDC or -8 VDC treatments were applied to the antennae. 100 µg of either (4a*S*,7*S*,7a*R*)-nepetalactone, (*R*)-linalool or (*E*)-β-farnesene was added to a piece of filter paper (10 µL applied from a 10 µg/µL diethyl ether solution) and placed for 30 min within a stream of charcoal-purified, humidified air flowing towards the antenna at 10 mL/min. Antennae were then dipped in 50 µL of re-distilled diethyl ether for 1 min to extract adsorbed compounds. Due to the low adsorption rates observed in preliminary tests, five antennae were extracted individually to create one replicate. Five replicates were collected per compound and voltage treatment. (4a*S*,7*S*,7a*R*)-Nepetalactone and (*E*)-β-farnesene were synthesised in house, assessing purity via NMR^52,53^, whilst (*R*)-linalool was purchased from Botanix Ltd. (Kent, England) and was 95% pure.

##### Gas-chromatography

Antennal extracts, injected in 4 µL aliquots, were analysed on an Agilent 8890 GC fitted with a non-polar HP1 column (50 m length × 0.32 mm inner diameter × 0.52 μm film thickness; J&W Scientific), using the following temperature programme: 30°C for 5 min, rising at 5°C/min to 150°C, followed by a 10°C/min rise to 230°C for a total run time of 60 min. Peak IDs were confirmed by GC peak enhancement via co-injection with authentic standards for (*R*)-linalool, (4a*S*,7*S*,7a*R*)-nepetalactone and (*E*)-β-farnesene^54^. Peak ID was further confirmed by comparison of mass spectra of extract peaks with authentic standards on an Agilent 5977B GC-MSD, using the same GC conditions as above, with ionisation by electron impact (70 eV, 220°C). Tentative identification of compounds was achieved by comparison of spectra with the National Institute of Standards and Technology (NIST) mass spectral library (2020, NIST, Gaithersburg, MD, USA). (*R*)-Linalool, (4a*S*,7*S*,7a*R*)-nepetalactone and (*E*)-β-farnesene amounts (ng) in antennal extracts were estimated using peak areas from a calibration curve of the respective authentic standard at 0.1, 1, 5 and 10 ng and generating a line of best fit equation (polynomial) in Microsoft Excel.

#### Antennal electrophysiology (electroantennography/EAG)

##### Baseline EAG recordings

Electrophysiological responses were recorded for test compounds on *A. fabae,* female *A. ervi, B. terrestris* and *D. melanogaster* antennae at doses of 0.1 ng, 1 ng, 10 ng, 100 ng, 1 µg, 10 µg and 100 µg (delivered in 10 µL solutions), using 10 µL redistilled diethyl ether as solvent control. EAG was performed as described previously^55^, with amendments. An antenna was carefully excised from a live insect and suspended between two electrodes made from Ag-AgCl borosilicate glass filled with Ringer solution (without glucose) and connected to silver wire (Ø 0.37 mm, Biochrom Ltd., Cambridge, UK). The base of the antenna was connected to a grounded electrode. A glass tube positioned approximately 5 mm away from the antennal preparation was connected to a stimulus controller (CS-02; Ockenfels Syntech GmbH, Kirchzarten, Germany) and facilitated a continuous flow of charcoal-purified humidified air towards the antenna at a rate of 1 L/min. The signal was passed through a high-impedance amplifier (UN-06, Syntech) and recorded using the Syntech EAG software package EAG v1.0 (6/1993). The absolute negative amplitude changes in response to the stimuli were recorded in mV and normalized against the positive controls (=100%), resulting in test stimuli being expressed as percentages^56^. Test compounds included (*E*)-β-farnesene, (*R*)-linalool, (*E*)-2-heptenal (Fluka, Germany, 98%), with (*E*)-β-caryophyllene (SAFC, St. Louis, MO, USA, ≥80%) as positive control for *A. fabae*; (*E*)-β-farnesene, (*R*)-linalool, (4a*S*,7*S*,7a*R*)-nepetalactone, with (*E*)-β-caryophyllene as positive control for *A. ervi*; (*E*)-ocimene (synthesized in house and assessed for purity via NMR)^57^, (*R*)-linalool, citral (Fluka, Germany, 95%), with benzaldehyde (Sigma-Aldrich, St. Louis, MO, USA, >99%) as positive control for *B. terrestris*; and (*R*)-limonene (Fluka, Germany, 98%), (*R*)-linalool, (*E*)-2-hexenal (SAFC, St. Louis, MO, USA, >95%), with (+)-fenchone (Fluka, Germany, 97%) as positive control for *D. melanogaster*. Ten replicates per test compound/species were recorded.

##### EAG recordings from antennae with a reduced state of charge (‘neutralisation’)

A modified setup was used to assess the effect of reducing the amount of antennal charge on EAG responses for *A. ervi* and *B. terrestris,* using 1 µg and 10 µg of (*R*)-linalool, respectively. Antennae were set up as described for baseline EAG recordings. On a single antenna, EAG responses were measured to i) 10 µL diethyl ether, ii) (*R*)-linalool and iii) (*R*)-linalool after the application of plasma bursts 5 mm from the antenna eight times. Recordings were repeated 24 times for *A. ervi* and 11 times for *B. terrestris,* leaving a 40-60 s lapse between stimulations. EAG responses were not normalised to a positive control due to the unknown effect of exposure to plasma on subsequent EAG responses.

##### EAG recordings from externally charged antennae

A modified EAG setup was used to assess the effect of applied charge on EAG responses across *A. fabae, A. ervi, B. terrestris* and *D. melanogaster*. Following the suspension of an insect antenna between two glass electrodes, a tungsten electrode (treatment electrode) connected to a function generator was brought into contact with the surface of the antenna from behind without obstructing the flow of VOCs toward the antennae. The treatment electrode was used to deliver a voltage bias onto the antennae at 0 V and ± 0.5, 1, 2, 4 and 8 VDC, using MATLAB R2022a to control for and visualise the applied voltage. EAG responses were recorded for the above-mentioned test compounds against their respective insect species, at a single dose, across increasing charges. The charge treatment was applied in random order. All charges were tested on a single antenna leaving a 40-60 s lapse between stimulations, with a minimum of seven replicates/compound/species. At the start and end of each replicate, EAG responses were recorded for the positive control and diethyl ether at 0 V and normalised to the positive control. The dose and replicate number for each compound/species tested are shown in table S4. Doses were chosen as the lowest dose required to induce a significant EAG response in baseline EAG recordings, and their respective ten-fold lower dose.

An extended version of the above experiment was done on *A. ervi* antennae against 100 ng (*R*)-linalool under 0 V and +4, 6, 8, 10 and 12 VDC stimulations to assess the effect of increased positive charge on EAG responses (n=12).

##### EAG dose-response recordings from externally charged antennae

The charged EAG experimental setup was used as described above, with some modifications. A -8 VDC potential was applied to a single antenna using a tungsten electrode as above. EAG responses to increasing doses of test compounds were sequentially recorded at 0.1 ng, 1 ng, 10 ng, 100 ng, 1 µg, 10 µg and 100 µg. Positive control and 10 µL diethyl ether at 0 V were run at the start and end of each replicate as described previously, with all data normalised to the positive control. Replication number varied depending on compound and species as follows: *A. fabae –* (*E*)-β-farnesene (n=11), (*R*)-linalool (n=8), (*E*)-2-heptenal (n=8); *A. ervi* - (*E*)-β-farnesene (n=7), (*R*)-linalool (n=5), (4a*S*,7*S*,7a*R*)-nepetalactone (n=7); *B. terrestris* - (*E*)-ocimene (n=6), (*R*)-linalool (n=6), citral (n=5) and *D. melanogaster* – (*R*)-limonene (n=5), (*R*)-linalool (n=5), (*E*)-2-hexenal (n=5).

#### Finite element modelling

A finite element model (FEM) was produced to-explore the biophysics-underlaying antennal boundary layer behaviour and electrostatics. We modelled the capture of minimally charged VOCs along an electrically biased antenna to assess the comparative influence of advection-diffusion and electrostatic forces. As measurables, product concentration and final deposition were evaluated. COMSOL Multiphysics software V 6.2 (COMSOL inc., Stockholm, Sweden) was used for this analysis. Due to the independence of the fluid and electrical processes, each were solved individually. Upon solving, the resulting fluid and electrical fields were used to solve the advection-diffusion-charge migration of a dilute suspension of VOC.

##### Antennal geometries

Two bio-inspired antenna morphologies were modelled based on SEMs and empirical data to ensure biologically relevant insight. One was formed with a dense canopy of hairs, like the honeybee and parasitic wasp antennae, and the other without such hairs, like the antenna of the black bean aphid. Together these models enable the assessment of how different morphological features affect olfactory capture and whether different forms show increased capture due to either fluid flow or electrostatics.

We studied both longitudinal fluid flows, parallel to the antenna, and crossflows, perpendicular to the antenna. For longitudinal sensing, the modelled geometry consisted of a 1.35 mm long cylindrical section of 0.1 mm radius with a spherical cap in a large surrounding domain (Figure 7A). Due to rotational symmetry, only one quarter of the structure was modelled using symmetry conditions in the x-y and x-z planes. The enclosing boundaries were placed at 10 mm, 10 mm and 10 mm distances from the antenna in x, y, z directions, respectively, and were thus 100 times the radius or 10 times the antennal length from the structure. Hairs of length 19 μm and radius 1.5 μm were placed in dense canopy with an offset configuration. The first row had an angular spacing of 10° over the 90° section of antenna, followed by a row with hairs offset by 5° and rows spaced by 0.01 mm. For longitudinal flow, hairs were placed over the spherical cap and in 30 rows along the first 0.3 mm section of the cylindrical form (Figure 7a). The domain was discretised using a tetrahedral mesh consisting of 3,261,274 boundary elements in the dense array case and 172,731 boundary elements for the hairless case. The large difference in mesh elements reflects the multi-scale nature of the dense hair problem, where the hair tips presented small thin regions that required a finer discretisation.

**Figure 7.**
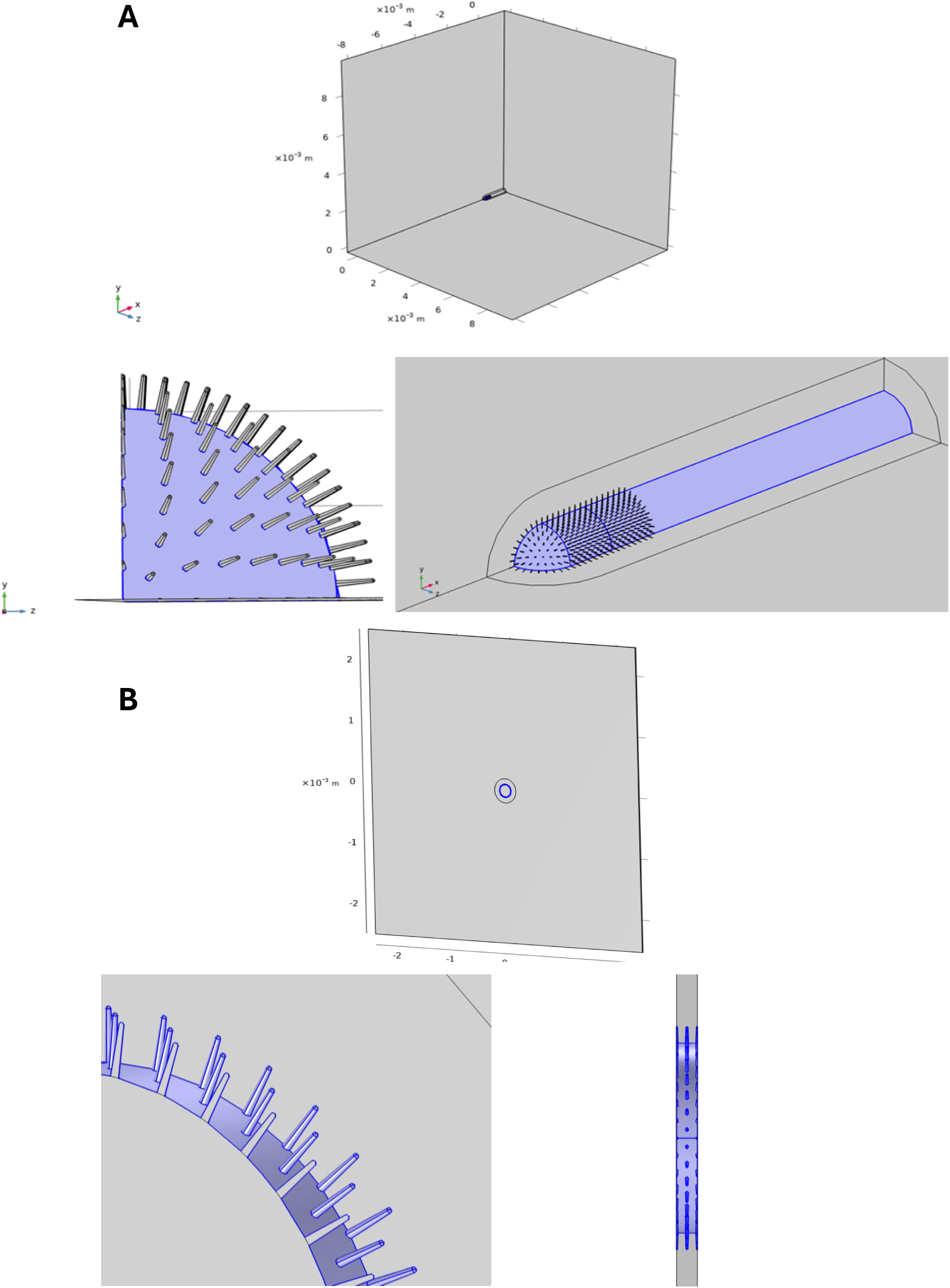
Geometry of modelled antenna and the computational domain. (A) Parallel flow: The far-field walls are placed at distances of 100 times the radius from the antenna to remove boundary effects. The antenna consists of a 1.35 mm long section with 0.1 mm radius, with hairs placed over the spherical cap and along the first 0.3 mm section of the cylindrical form. The hair lengths are 0.01 mm. (B) Perpendicular flow: The far-field walls are placed at distances of 50 times the radius from the antenna to remove boundary effects. The antenna consists of a 0.02 mm long section with 0.1 mm radius. The hair lengths are 0.01 mm.

For crossflow, we modelled a 0.2 mm short cross-section of the antenna aligned with the x-axis, using symmetry conditions at two parallel y-z planes 0.02 mm apart. Only two-rows of hairs were required to obtain results for an effectively infinite antenna. The same antennal geometry was otherwise considered (Figure 7B). The enclosing boundaries were placed at 5 mm, 5 mm and 0.02 mm in x, y, z directions, respectively. The domain was discretised using a tetrahedral mesh to solve the equations consisting of 3,033,113 boundary elements in the dense array case and 295,254 boundary elements for the hairless case.

In both cases, a mesh independence study was carried out to ensure accuracy of the computed results. Considering the most sensitive case, a dense hair array and 𝑈_∞_ = 0.1 𝑚𝑠^−1^, the following results were obtained for finer meshes:

##### Fluid-antenna interaction modelling

The steady interaction between a uniform flow of air with a constant fluid density and a fixed antenna within the described computational domain is computed by the incompressible Navier-Stokes equations:

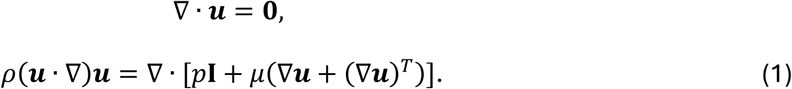

These equations are solved on a fixed mesh subject to boundary conditions (given below), denoting the three-dimensional fluid velocity by 𝒖 (m/s), the dynamic viscosity by 𝜇 (Ns/m^2^), the pressure field by p (kg /m s^2^) and the fluid density by 𝜌 (kg/m^3^). Since the background fluid is air, we set the dynamic viscosity to be 1.81x10^-5^ Ns/m^2^ and fluid density to be 1 kg/m^3^ at 293K.

Boundaries occur at the edge of the domain and on the antenna. Conditions were prescribed therein to ensure physically accurate and consistent results. At the inlet, the direction from which the flow comes upstream of the antenna, the far-field flow was prescribed as 𝑢 = 𝑈_∞_, 𝑣 = 0, 𝑤 = 0, where 𝑢, 𝑣, 𝑤 denote the velocity of the fluid in Cartesian directions 𝑥, 𝑦, 𝑧, and 𝑈_∞_ denotes the magnitude of the freestream flow speed far from the antenna. To simulate a range of appropriate flight speeds for an insect, we evaluated three scenarios with 𝑈_∞_ = 0.001, 0.01 and 0.1 m/s. The outlet boundary condition, downstream of the antenna in the x-direction, was prescribed to be 𝑝 = 0. Along the antenna, a no-slip wall condition was applied 𝑢 = 0, 𝑣 = 0, 𝑤 = 0, leading to a boundary layer along the antenna. A slip wall condition was applied to the upper x-z boundary of the domain to constrain the flow with: 𝑢 = 𝑈_∞_, 𝑣 = 0, 𝑤 = 0. Symmetry conditions were applied, as previously stated, in the bounding x-y and x-z planes of the geometry due to the rotationally symmetric nature of the geometry.

##### Computing the electrostatic field

The electrostatic field throughout the domain was governed by the equations:

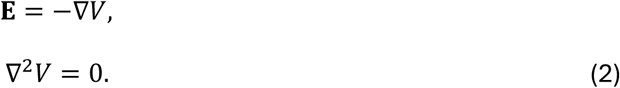

Here, V denotes the surface electric potential and 𝐄 the electric field. A –8 V potential was applied to the antenna surface based on the bias applied during the experiments. Within the bounding domain, symmetric conditions were applied as above, and all other boundaries were set to 0 V far from the antenna. Regarding the hairs, their individual charge or potential were not experimentally measured and thus not prescribed here. The hairs are treated as a dielectric with a relative permittivity of 14 that polarize in the presence of the biased cuticle.

##### Modelling the transport of VOCs in the domain

The transport of the dilute substance in the domain was calculated by the following equation:

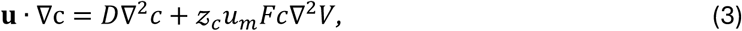

where 𝒖 is given by (1) and 𝑉 by (2) to determine the comparative effect of electrostatic forces and fluid dynamic influences in the transport and capture of VOCs. In (3), 𝑐 indicates the concentration of the substance in air (mol/m^3^), 𝐷 is the diffusion coefficient, which was defined as 6.7x10^-7^m^2^/s and 𝓏_𝑐_ is the charge number of the substance (dimensionless) given in the units of elementary charge to quantify the charge of ions or single molecules. We consider this to represent an “effective charge”, since the dipole moment of VOCs produce forces several orders of magnitude below that related to an elementary charge. Hence, we considered values of 𝓏_𝑐_ = 0.00001, 0.0001, 0.001, 0.1, 1 to show the broad range and influence of VOC charge from weakly polarized dipoles to an ionic molecule of one elementary charge. The ionic mobility, 𝑢_𝑚_ = D/RT, is also calculated from input parameters, whereby R is the molar gas constant (J/mol·K) and T = 293 is temperature (K). Finally, F denotes Faraday’s constant (A·s/mol). The diffusion coefficient of 5x10^-6^ m^2^/s was chosen as a representative value across molecules which may underestimate VOC catch results for molecules with a large diffusion coefficient. However, this model aims to show the relative effects of electrostatics on VOC olfactory capture and therefore the use of exact diffusion coefficients for each molecule is not required.

For the boundary condition along the antennal surface and hairs, we set 𝑐 = 0 to simulate absorption of VOCs. The inflow concentration was 𝑐_0_ = 1 mol/m^3^ at the inlet boundary to model a uniform well-mixed distribution of VOCs in the oncoming flow. Symmetry conditions were again applied to the relevant boundaries. Outlet conditions are applied to all other boundaries with 𝐧 · 𝐷∇𝑐 = 0, 𝐧 the local normal of the surface. Our metric of interest here is a modified version of that presented in Claverie et al. (2022)^4^, and is denoted as the capture rate of the antenna given by:

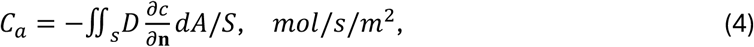

where S is the surface area of the modelled antennal section (including the hairs when present). From (4), the local gradient of the concentration over the antennal surface gives the capture rate. We divide the integrated value by the modelled surface area to enable comparisons between scenarios, since the antennal surface is much larger in the longitudinal case and when hairs are present (Table 3).

**Table 3:** Capture rates, 𝐶_𝑎_, mol/s/m^2^, of an antenna with a dense hair coverage and no hairs for different flow speeds and morphology. There is a monotonic trend in capture rate with the effective charge of the VOCs, which is consistent across flow speeds. However, when the charge number is at least 0.1, the capture rate becomes invariant to the flow speed, indicating that the electrostatic contribution to olfactory capture dominates transport forces due to the fluid flow.

Longitudinal flow
| Charge Number (q) | U = 0.001 m/s |  | U = 0.01 m/s |  | U = 0.1 m/s |  |
| --- | --- | --- | --- | --- | --- | --- |
|  | Dense | None | Dense | None | Dense | None |
| 0 | 0.0193 | 0.0276 | 0.0241 | 0.0344 | 0.0393 | 0.0558 |
| 0.00001 | 0.0194 | 0.0277 | 0.0242 | 0.0345 | 0.0393 | 0.0558 |
| 0.0001 | 0.0196 | 0.0280 | 0.0244 | 0.0349 | 0.0395 | 0.0562 |
| 0.001 | 0.0223 | 0.0320 | 0.0271 | 0.0388 | 0.0422 | 0.0602 |
| 0.01 | 0.0584 | 0.0867 | 0.0616 | 0.0913 | 0.0739 | 0.1087 |
| 0.1 | 0.5413 | 0.8192 | 0.5414 | 0.8193 | 0.5416 | 0.8198 |
| 1 | 5.3954 | 8.1641 | 5.4115 | 8.1889 | 5.4109 | 8.1878 |

Crossflow
| Charge Number (q) | U = 0.001 m/s |  | U = 0.01 m/s |  | U = 0.1 m/s |  |
| --- | --- | --- | --- | --- | --- | --- |
|  | Dense | None | Dense | None | Dense | None |
| 0 | 0.0068 | 0.0110 | 0.0111 | 0.0177 | 0.0217 | 0.0347 |
| 0.00001 | 0.0068 | 0.0110 | 0.0111 | 0.0178 | 0.0217 | 0.0347 |
| 0.0001 | 0.0069 | 0.0112 | 0.0112 | 0.0180 | 0.0218 | 0.0349 |
| 0.001 | 0.0081 | 0.0132 | 0.0126 | 0.0202 | 0.0231 | 0.0371 |
| 0.01 | 0.0281 | 0.0461 | 0.0314 | 0.0511 | 0.0391 | 0.0634 |
| 0.1 | 0.2863 | 0.4729 | 0.2906 | 0.4752 | 0.2913 | 0.4758 |
| 1 | 2.2795 | 4.0892 | 2.7642 | 4.5853 | 2.8880 | 4.7412 |

### Quantification and statistical analysis

For the statistical analysis of faraday cup measurements, raw charge recordings were converted to picocoulomb (pC) measurements using MATLAB. All statistical analyses were carried out in RStudio V2023.12.1+402 running R V4.3.3^58^. All data within species were tested for normality and outliers removed using the “stats” V.4.3.3 package. Kruskal-Wallis and Dunn post-hoc tests, packages “stats” V4.3.3 and “dunn.test” V1.3.6 respectively, were carried out to compare charge measurements between all treatments within both *B. terrestris* and *A. fabae*. For *A. ervi,* this was carried out using Anova and Tukey post-hoc tests, package “stats” V4.3.3. Data were visualised in R.

Within each compound treatment, the amount of compound in antennal extracts was compared across charge treatments (0 V, +8 VDC and -8 VDC). The data were tested for normality via Shapiro-Wilk test. ANOVA and Tukey post-hoc tests were used for (4a*S*,7*S*,7a*R*)-nepetalactone and (*R*)-linalool, whilst Kruskal-Wallis and Dunn post-hoc tests were used for (*E*)-β-farnesene.

For baseline EAG experiments and charged dose-response recordings, normalised EAG responses were tested for normality by Shapiro-wilk test and either a Student’s t-test or Wilcoxon test (depending on normality, “Stats” package V4.3.3) was used to compare responses between each compound dose and diethyl ether.

For neutralised EAG recordings, within-species data were tested for normality using Shapiro-wilk test. For *A. ervi*, EAG responses for diethyl ether and (*R*)-linalool before and after neutralisation were statistically compared by Kruskal Wallis (“Stats” package V4.3.3) and Dunn post-hoc tests (“dunn.test” V1.3.6). For *B. terrestris*, Kruskal Wallis and Wilcoxon post-hoc test (“rstatix” package V 0.7.2) were used.

For charged EAG experiments, EAG recordings from within species and per compound were statistically compared between each charge treatment and its respective diethyl ether control. Data were tested for normality via Shapiro-wilk test. Depending on data distribution, either Student’s t-tests or Wilcoxon test (“Stats” package V4.3.3) was carried out between EAG responses at each charge treatment and the diethyl ether control. The statistical test used and respective P-values are described in Table S2.

Correlation analysis was carried out between normalised EAG response, voltage treatment and antennal dimensions. Excised antennae across all four species were imaged under light microscopy (Zeis Axioimager Z2 with Zeis Axiocam 512 colour camera, Supplementary 2) and antennal length, antennal inner and outer diameter and cuticle thickness measurements were taken. Antennal surface area was calculated for each species under the assumption that the antenna is cylindrical. Since data were not normally distributed, a Q-Q plot (“car” package V.3.1.2) was made to assess the most appropriate generalised linear model (GLM) distribution that fits the data. An Akaike information criterion (AIC) test was used to assess which GLM distribution between gamma, inverse gaussian or Tweedie distribution best fits the data (“Stats” package V4.3.3). A GLM with inversion Gaussian distribution was used to compare the interaction between normalised EAG response, charge treatment and antennal surface area (“Stats” package V4.3.3). Bootstrap analysis was carried out to assess accuracy for GLM outputs (“boot” package V1.3.30). Datapoints were visualised via scatterplot. Correlation analysis was carried out as above within each species to compare the relationship/interaction between normalised EAG response, voltage treatment and compound dipole moment.

All statistical analyses and visualization were carried out in R V4.3.3.

## Supplemental Information

Supplementary 1 - Supplementary figures and tables. (.doc) Supplementary 2 – Bioimaging methods (.doc)

