## Supplemental tables and figures for "Electrostatic facilitation of odorant capture in insects"

Supplementary information for: Electrostatic facilitation of odorant capture in insects.

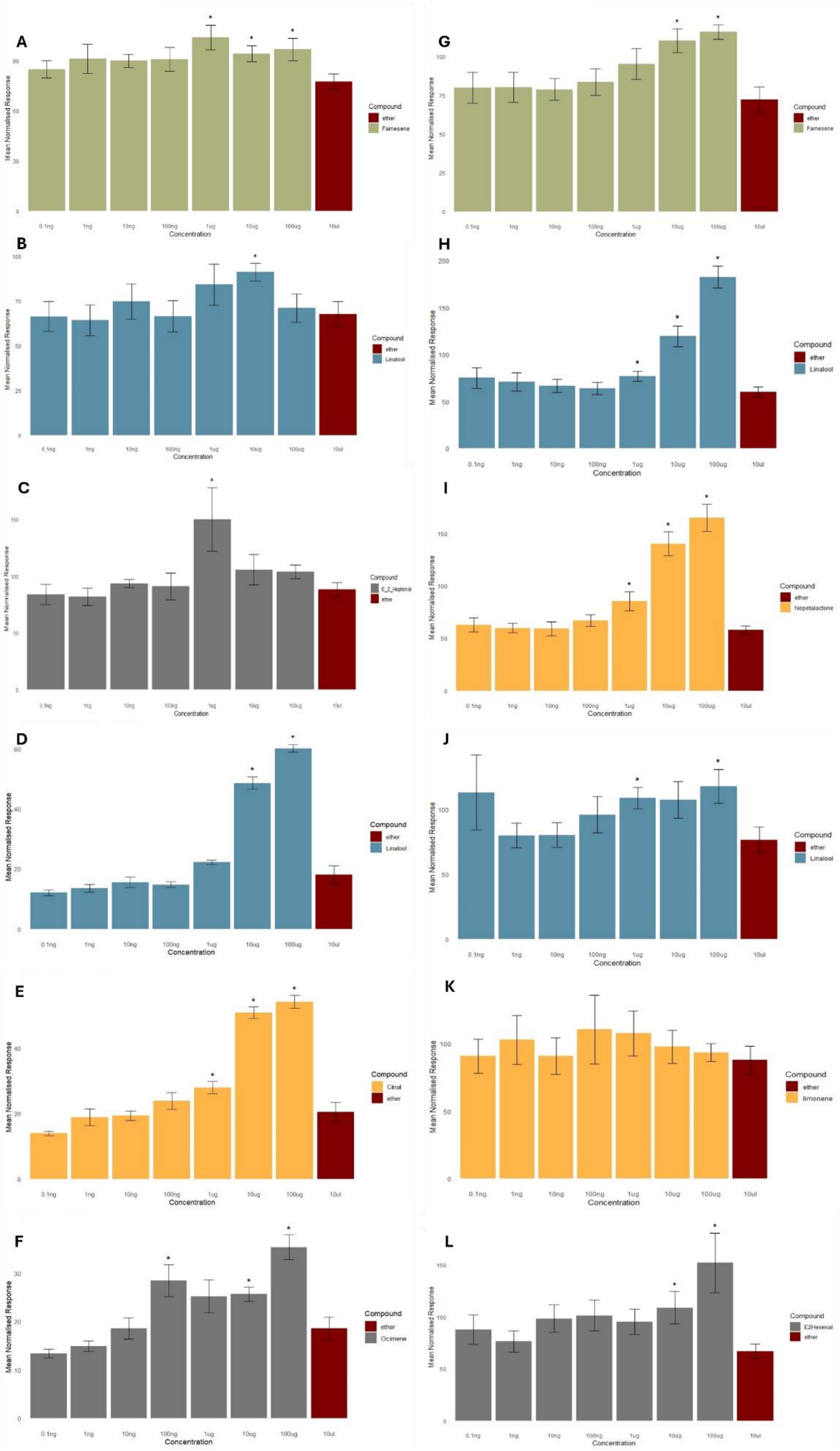

**Figure S1**: Electrophysioligical (EAG) responses to increasing concentrations of test compounds from antennae of; *Aphis fabae* against A: (*E*)-β-farnesene, B: (R)-linalool and C: (*E*)-2-heptenal (n=8), *Bombus terrestis* against D: (R)-linalool, E: Citral and F: (*E*)-ocimene, *Aphidius ervi* against G: (*E*)-β-farnesene, H: (R)-linalool and I: (4aS,7S,7aR)-nepetalactone and *Drosophila melanogaster* against J: (R)-linalool, K: Limonene and L: (*E*)-2-hexenal. EAG responses were normalised to a positive control: *B. terrestris* = benzaldehyde, *A. fabae* and *A. ervi* = (*E*)-caryophyllene and *D. melanogaster* = (+)-fenchone. Ten replicates were recorded per test compound and insect. Significance from diethyl ether solvent control: ·=p<0.1, *=p<0.05.

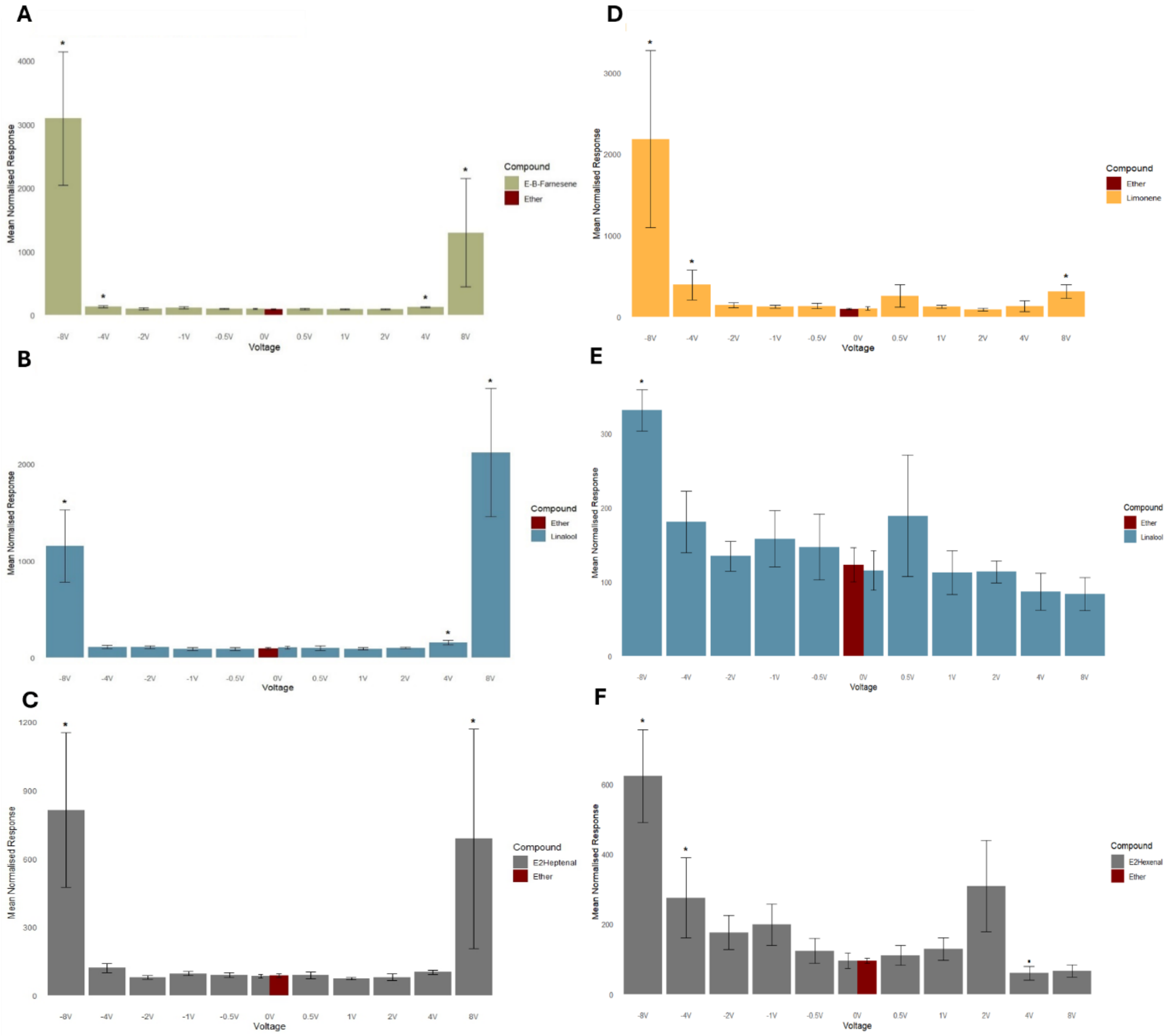


**Figure S2:** Electrophysiological (EAG) responses of *Aphis fabae* and *Drosophila melanogaster* antennae, exposed to a range of voltages, to synthetic compounds at a dose 10-fold lower than a significant EAG-active dose (baseline measurements Figure S1, mean ±SE). A: *A. fabae*, (E)-β-farnesene (n=8, dose=100ng), B: *A. fabae*, (R)-linalool (n=8, dose=1μg), C: *A. fabae*, (*E*)-2-heptenal (n=8, dose=100 ng), D: *D. melanogaster,* Limonene (n=7, dose=100 μg), E: *D. melanogaster,* (R)-linalool (n=8, dose=100 ng) and F: *D. melanogaster*, (*E*)-2-heptenal (n=7, dose=1 μg). EAG responses were normalised to a positive control: *A. fabae* = (E)-caryophyllene, *D. melanogaster* = (+)-fenchone. Charge was applied on antennae via tungsten electrode. Significance from diethyl ether solvent control: ·=p<0.1, *=p<0.05.

**Table S1** Difference in electrophysiological (EAG) responses from *A. fabae*, *B. terrestris, D. melanogaster* and *A. ervi* antennae between an applied voltage (V) and control (diethyl ether), representing the effect of applied charge on EAG responses, represented as a % to control.

| Voltage (V) | Mean normalised response (%) | Response change Vs Ether control (%) |
|---|---|---|
| | *Aphis fabae* (aphid) | |
| | (*E*)-2-heptenal | |
| *8* | 687.4 ±481 | 744.4 |
| *4* | 101.9 ±8.9 | 110.3 |
| *2* | 79.9 ±14.3 | 86.5 |
| *1* | 73.3 ±5.9 | 79.3 |
| *0.5* | 88.5 ±14.7 | 95.8 |
| *0* | 84.1 ±6.8 | 91.1 |
| *-0.5* | 89.3 ±9.5 | 96.7 |
| *-1* | 95 ±9.3 | 102.9 |
| *-2* | 79.3 ±7.2 | 85.8 |
| *-4* | 119.9 ±19.2 | 129.8 |
| *-8* | 813.3 ±339.3 | 880.8 |
| | (*E*)-β-farnesene | |
| *8* | 1298 ±849.9 | 1405.8 |
| *4* | 126 ±10 | 136.5 |
| *2* | 96.1 ±8.6 | 104.1 |
| *1* | 91.8 ±6.5 | 99.4 |
| *0.5* | 99.8 ±12 | 108.0 |
| *0* | 100.8 ±9.3 | 109.1 |
| *-0.5* | 102.3 ±7.4 | 110.7 |
| *-1* | 118.5 ±15.7 | 128.3 |
| *-2* | 102.5 ±15.6 | 111.0 |
| *-4* | 137.8 ±17.1 | 149.2 |
| *-8* | 3094.4 ±1047.9 | 3351.3 |
| | Linalool | |
| *8* | 2117.9 ±662.4 | 2293.7 |
| *4* | 156.1 ±23.2 | 169.1 |
| *2* | 102.5 ±8 | 111.0 |
| *1* | 92.4 ±13.2 | 100.0 |
| *0.5* | 100.1 ±25.9 | 108.4 |
| *0* | 104.5 ±13.6 | 113.2 |
| *-0.5* | 90.1 ±12.5 | 97.6 |
| *-1* | 91.4 ±12.5 | 99.0 |
| *-2* | 108 ±14.1 | 117.0 |
| *-4* | 111.5 ±15.1 | 120.8 |
| *-8* | 1156 ±373.8 | 1252.0 |
| | Ether | |
| *0* | 92.3 ±5 | 100.0 |
| | *Bombus terrestris* (bumble bee) | |
| | Citral | |
| *8* | 64.7 ±13.3 | 158.0 |
| *4* | 34.6 ±4.8 | 84.4 |
| *2* | 36.4 ±4 | 89.0 |
| *1* | 38.7 ±5.9 | 94.5 |
| *0.5* | 59.4 ±19.3 | 145.1 |
| *0* | 46 ±3.2 | 112.3 |
| *-0.5* | 76.3 ±20.8 | 186.3 |
| *-1* | 116 ±62.7 | 283.3 |
| *-2* | 110.4 ±59.1 | 269.7 |
| *-4* | 198.1 ±108.1 | 483.8 |
| *-8* | 274.6 ±165.1 | 670.5 |
| | (*E*)-ocimene | |
| *8* | 39 ±10.6 | 95.2 |
| *4* | 30.9 ±5.9 | 75.3 |
| *2* | 33.4 ±3.4 | 81.6 |
| *1* | 37.1 ±3.7 | 90.7 |
| *0.5* | 45.7 ±6.2 | 111.6 |
| *0* | 39 ±4.6 | 95.2 |
| *-0.5* | 53.4 ±6.6 | 130.5 |
| *-1* | 49.3 ±7 | 120.3 |
| *-2* | 61.3 ±12.3 | 149.7 |

| | | |
|---|---|---|
| *-4* | 69.6 ±12.4 | 169.9 |
| *-8* | 144 ±38.3 | 351.6 |
| Linalool | | |
| *8* | 101 ±31.7 | 246.6 |
| *4* | 32.1 ±4.4 | 78.5 |
| *2* | 32.6 ±2.4 | 79.5 |
| *1* | 45 ±3.9 | 109.9 |
| *0.5* | 41.9 ±6.1 | 102.2 |
| *0* | 45.6 ±4.8 | 111.3 |
| *-0.5* | 59.4 ±7.2 | 145.1 |
| *-1* | 61.3 ±8.4 | 149.7 |
| *-2* | 89 ±19.7 | 217.3 |
| *-4* | 134.7 ±37.2 | 329.0 |
| *-8* | 191.4 ±27.2 | 467.4 |
| Ether | | |
| *0* | 41 ±2.7 | 100.0 |
| *Drosophila melanogaster* (fruit fly) | | |
| (*E*)-2-hexenal | | |
| *8* | 66.7 ±17.1 | 62.5 |
| *4* | 60 ±19.6 | 56.2 |
| *2* | 308.7 ±130.2 | 289.3 |
| *1* | 129.7 ±32.1 | 121.5 |
| *0.5* | 110.7 ±28.8 | 103.7 |
| *0* | 95.4 ±22.1 | 89.4 |
| *-0.5* | 123.4 ±35.5 | 115.6 |
| *-1* | 198.6 ±58.8 | 186.1 |
| *-2* | 176.1 ±48.6 | 165.0 |
| *-4* | 275.3 ±114.1 | 257.9 |
| *-8* | 622.7 ±133 | 583.5 |
| Limonene | | |
| *8* | 313.3 ±84.6 | 293.5 |
| *4* | 135.1 ±66.8 | 126.6 |
| *2* | 92.1 ±16.9 | 86.3 |
| *1* | 128.3 ±16.8 | 120.2 |
| *0.5* | 260.1 ±137.8 | 243.7 |
| *0* | 105.4 ±20.3 | 98.8 |
| *-0.5* | 136.4 ±33 | 127.8 |
| *-1* | 127.1 ±17 | 119.1 |
| *-2* | 145.3 ±30.5 | 136.1 |
| *-4* | 395.9 ±183.8 | 370.9 |
| *-8* | 2184.6 ±1087.9 | 2046.9 |
| Linalool | | |
| *8* | 83.6 ±22.4 | 78.4 |
| *4* | 87 ±25 | 81.5 |
| *2* | 113.6 ±14.8 | 106.5 |
| *1* | 112.5 ±29.8 | 105.4 |
| *0.5* | 189.1 ±81.6 | 177.2 |
| *0* | 115.4 ±26.7 | 108.1 |
| *-0.5* | 147 ±44.2 | 137.7 |
| *-1* | 158.3 ±37.8 | 148.3 |
| *-2* | 134.6 ±20.3 | 126.1 |
| *-4* | 181 ±41.2 | 169.6 |
| *-8* | 331.3 ±27.8 | 310.4 |
| Ether | | |
| *0* | 106.7 ±9.1 | 100.0 |
| *Aphidius ervi* (parasitoid wasp) | | |
| *(E)*-β-farnesene | | |
| *8* | 20 ±3.9 | 21.3 |
| *4* | 22.6 ±4.7 | 24.0 |
| *2* | 45 ±8.6 | 47.9 |
| *1* | 76.6 ±10 | 81.6 |
| *0.5* | 72.9 ±24.6 | 77.6 |
| *0* | 105.4 ±38 | 112.3 |
| *-0.5* | 82.6 ±21.7 | 88.0 |
| *-1* | 163.7 ±62.2 | 174.4 |
| *-2* | 154.4 ±29.9 | 164.5 |
| *-4* | 199.6 ±85.6 | 212.6 |

| | | |
|---|---|---|
| *-8* | 228 ±70.3 | 242.9 |
| Linalool | | |
| *8* | 51.8 ±17.7 | 55.1 |
| *4* | 23.4 ±6.2 | 24.9 |
| *2* | 74 ±37.3 | 78.8 |
| *1* | 51.4 ±10 | 54.7 |
| *0.5* | 84 ±21.4 | 89.5 |
| *0* | 85.1 ±23.6 | 90.7 |
| *-0.5* | 136.4 ±32.9 | 145.3 |
| *-1* | 141 ±47.5 | 150.2 |
| *-2* | 152.1 ±20.6 | 162.1 |
| *-4* | 305.9 ±79.7 | 325.9 |
| *-8* | 327.3 ±104.5 | 348.7 |
| Nepetalactone | | |
| *8* | 193.5 ±65.3 | 206.2 |
| *4* | 80.7 ±14.7 | 85.9 |
| *2* | 88.9 ±7.7 | 94.7 |
| *1* | 85.3 ±13.3 | 90.9 |
| *0.5* | 143.7 ±19.3 | 153.1 |
| *0* | 81.2 ±12.1 | 86.5 |
| *-0.5* | 64.9 ±8.1 | 69.1 |
| *-1* | 86.2 ±10.9 | 91.8 |
| *-2* | 95.7 ±11.1 | 101.9 |
| *-4* | 136.8 ±25.5 | 145.8 |
| *-8* | 222.6 ±39.9 | 237.2 |
| Ether | | |
| *0* | 93.9 ±8.9 | 100.0 |

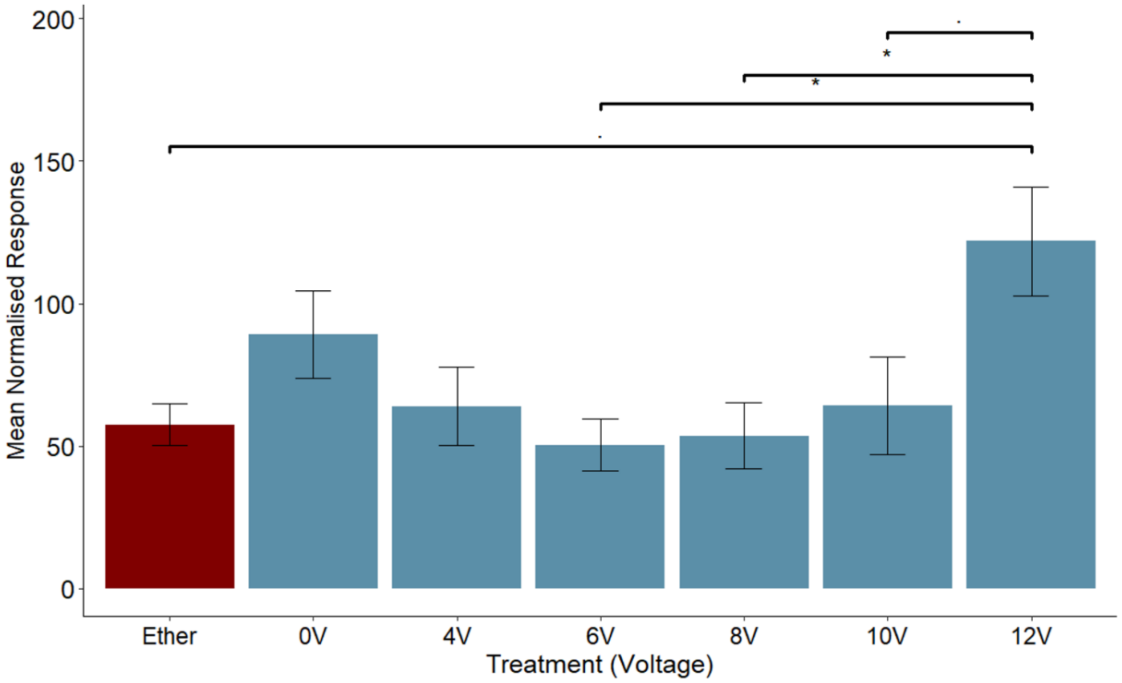


**Figure S3:** Electrophysiological (EAG) responses of *A. ervi* antennae to a 100ng dose of (R)-linalool exposed to a range of positive voltages (n=13). EAG responses were normalised to (*E*)-caryophyllene as a positive control. Charge was applied on antennae via tungsten electrode. Significance from diethyl ether solvent control: ·=p<0.1, *=p<0.05.

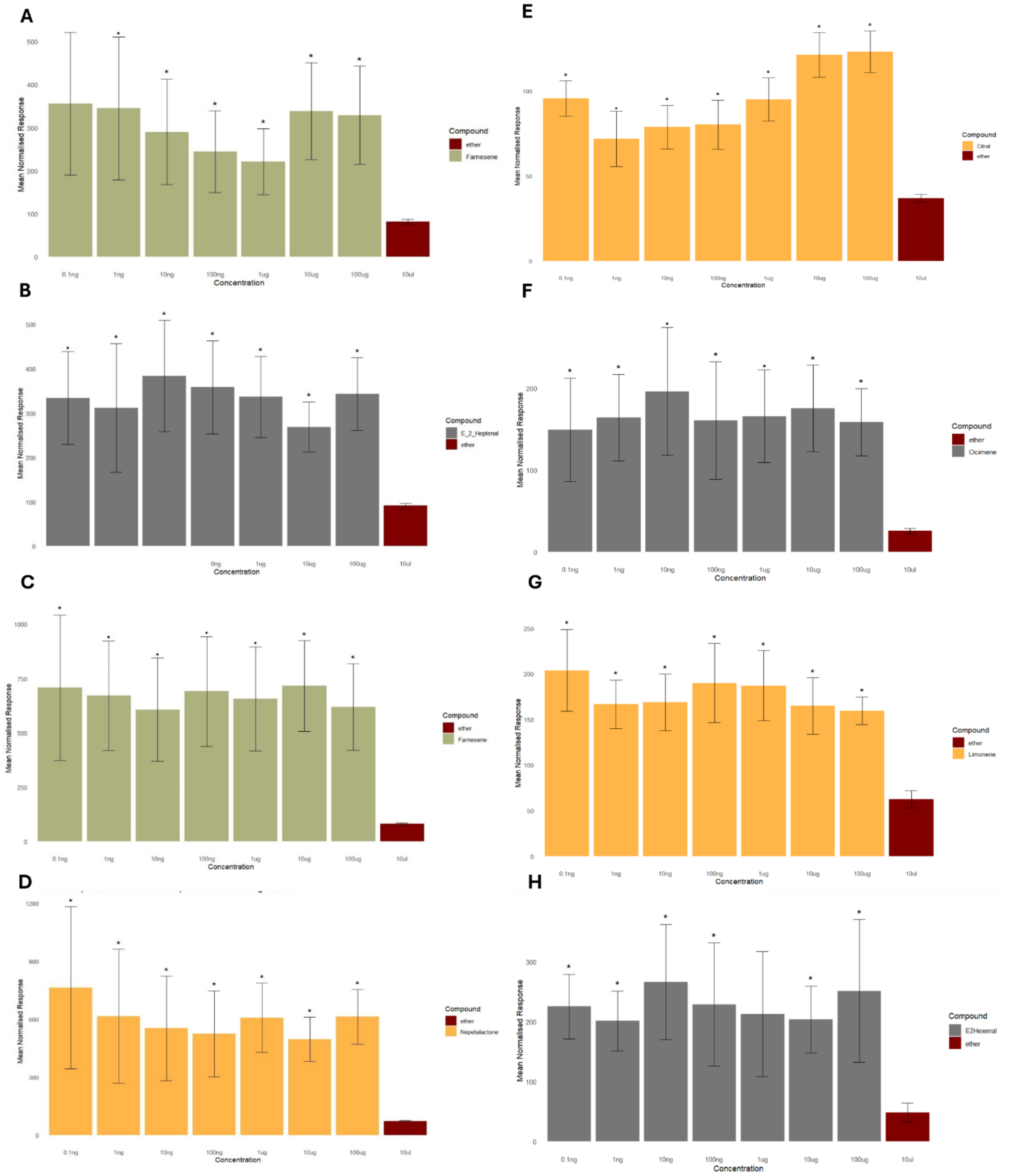


**Figure S4:** Electrophysiological (EAG) responses of *A. fabae, A. ervi, B. terrestris* and *D. melanogaster* antennae to a range of doses from synthetic compounds whilst biassed with -8 V (mean ±SE). A: *A.* fabae, (*E*)-β-farnesene (n=11), B: *A. fabae*, (*E*)-2-heptenal (n=8), C: *A. ervi*, (*E*)-β-farnesene (n=6), D: *A. ervi*, (4aS,7S,7aR)-nepetalactone (n=7), E: *B. terrestris*, Citral (n=5), F: *B. terrestris*, (*E*)-ocimene (n=6), G: *D. melanogaster*, limonene (n=5), H: *D. melanogaster*, (*E*)-2-hexenal (n=5). Significance from diethyl ether solvent control: ·=p<0.1, *=p<0.05.

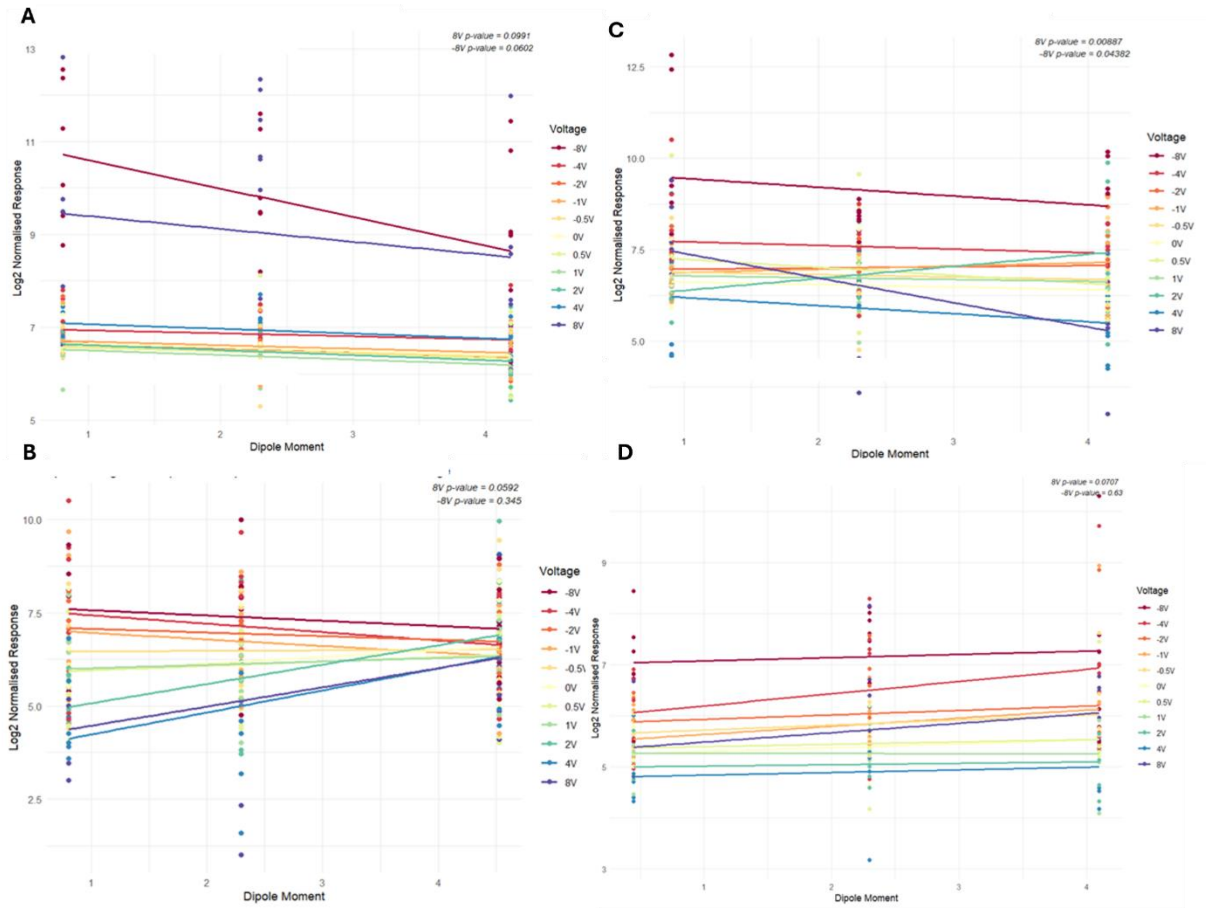


**Figure S5:** Relationship between electrophysiological (EAG) response and compound dipole moment from antennae of *A. fabae* (A), *A. ervi* (B), *B. terrestris* (C) and *D. melanogaster* (D) exposed to a range of voltages (V). EAG responses were taken from *B. terrestris* exposed to 1 µg (*R*)-linalool, 100 ng citral and 10 ng (*E*)-ocimene (total n=21), *A. ervi* exposed to 100 ng (*R*)-linalool, 100 ng (4aS,7S,7aR)-nepetalactone and 1 µg (*E*)-β-farnesene (total n=24), *A. fabae* exposed to 1 µg (*R*)-linalool, 100 ng (*E*)-2-heptenal and 100 ng (*E*)-β-farnesene (total n=24) and *D. melanogaster* exposed to 100 ng (*R*)-linalool, 100 µg (*R*)-limonene and 1 µg (*E*)-2-hexenal (total n=22). Significance compared to normalised EAG response interaction between +/- 8 V and 0 V, GLM.

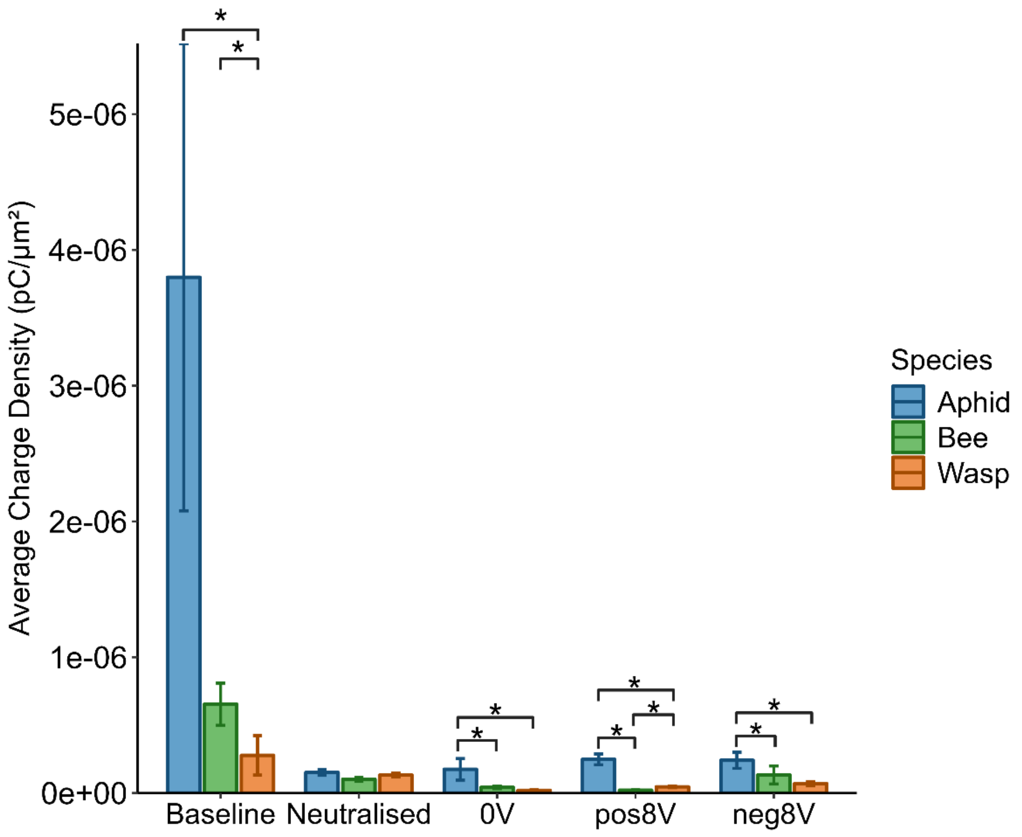


**Figure S6**: Average charge density (pC/µm$^2$) of *A. fabae*, *B. terrestris* and *A. ervi* antennae across baseline (no treatment), neutralised (exposed to a plasma beam to reduce spatial charge) or 0 V, +8 V and -8 V applied via tungsten electrode. Measurement of net charge on individual antennae was made with a Faraday cup (n=10 antennae/species, Figure 2). Antennal surface area was calculated from stereomicroscopy images (n=3 antennae/species). Significance: *=P<0.05, ANOVA/Tukey.

**Table S2**: Statistical test and P-values for electrophysiological (EAG) responses of *Aphis fabae, Aphidius ervi, Bombus terrestris* and *Drosophila melanogaster* antennae exposed to a range of voltages and synthetic compounds at a dose 10-fold lower than a significant EAG-active dose. This table supports data presented in Electrostatic facilitation of odorant capture in insects, Borg, *et al*. 2026, Figure 3 and S2.

| ***Compound*** | **Voltage** | **shapiro_p_value** | **test_name** | **p_value** |
|---|---|---|---|---|
| | | *Bee* | | |
| *Linalool* | 0V | 0.70254 | t-test | 0.421412 |
| *Linalool* | -2V | 0.189579 | t-test | 0.050638 |
| *Linalool* | 8V | 0.008921 | Wilcoxon test | 0.009281 |
| *Linalool* | 4V | 0.166332 | t-test | 0.115938 |
| *Linalool* | 0.5V | 0.056319 | t-test | 0.895346 |
| *Linalool* | -8V | 0.408939 | t-test | 0.001401 |
| *Linalool* | 2V | 0.810298 | t-test | 0.033032 |

| | | | | |
|---|---|---|---|---|
| *Linalool* | -0.5V | 0.162797 | t-test | 0.044645 |
| *Linalool* | 1V | 0.149135 | t-test | 0.413674 |
| *Linalool* | -4V | 0.468023 | t-test | 0.045281 |
| *Linalool* | -1V | 0.051613 | t-test | 0.054192 |
| *Citral* | 0V | 0.32521 | t-test | 0.250918 |
| *Citral* | -2V | 7.02E-05 | Wilcoxon test | 0.075228 |
| *Citral* | 8V | 0.492932 | t-test | 0.1261 |
| *Citral* | 4V | 0.322541 | t-test | 0.277303 |
| *Citral* | 0.5V | 4.52E-05 | Wilcoxon test | 0.3258 |
| *Citral* | -8V | 0.00012 | Wilcoxon test | 0.000909 |
| *Citral* | 2V | 0.686389 | t-test | 0.371978 |
| *Citral* | -0.5V | 0.000929 | Wilcoxon test | 0.007345 |
| *Citral* | 1V | 0.893677 | t-test | 0.737766 |
| *Citral* | -4V | 0.00012 | Wilcoxon test | 0.001451 |
| *Citral* | -1V | 8.99E-05 | Wilcoxon test | 0.066999 |
| *Ocimene* | -0.5V | 0.489706 | t-test | 0.117169 |
| *Ocimene* | -2V | 0.21793 | t-test | 0.15381 |
| *Ocimene* | 0.5V | 0.005536 | Wilcoxon test | 0.366559 |
| *Ocimene* | 8V | 0.000122 | Wilcoxon test | 0.055863 |
| *Ocimene* | 2V | 0.02017 | Wilcoxon test | 0.0944 |
| *Ocimene* | -8V | 0.186477 | t-test | 0.036118 |
| *Ocimene* | -1V | 0.319504 | t-test | 0.298156 |
| *Ocimene* | 1V | 0.818803 | t-test | 0.419012 |
| *Ocimene* | -4V | 0.556521 | t-test | 0.06122 |
| *Ocimene* | 0V | 0.01354 | Wilcoxon test | 0.595372 |
| *Ocimene* | 4V | 0.00971 | Wilcoxon test | 0.025695 |
| | *Wasp* | | | |
| *Linalool* | 0V | 0.21912 | t-test | 0.737045 |
| *Linalool* | -2V | 0.159418 | t-test | 0.027319 |
| *Linalool* | 8V | 0.199911 | t-test | 0.057645 |
| *Linalool* | 4V | 0.376304 | t-test | 2.68E-07 |
| *Linalool* | 0.5V | 0.118015 | t-test | 0.679485 |
| *Linalool* | -8V | 0.00702 | Wilcoxon test | 0.001585 |
| *Linalool* | 2V | 0.000569 | Wilcoxon test | 0.034666 |
| *Linalool* | -0.5V | 0.061804 | t-test | 0.247143 |

| | | | | |
|---|---|---|---|---|
| *Linalool* | 1V | 0.763908 | t-test | 0.004973 |
| *Linalool* | -4V | 0.062276 | t-test | 0.032543 |
| *Linalool* | -1V | 0.035436 | Wilcoxon test | 0.969643 |
| *E-B-Farnesene* | 0V | 0.011309 | Wilcoxon test | 0.413212 |
| *E-B-Farnesene* | -2V | 0.732613 | t-test | 0.086695 |
| *E-B-Farnesene* | 8V | 0.336574 | t-test | 1.12E-08 |
| *E-B-Farnesene* | 4V | 0.011053 | Wilcoxon test | 0.000148 |
| *E-B-Farnesene* | 0.5V | 0.036004 | Wilcoxon test | 0.154778 |
| *E-B-Farnesene* | -8V | 0.144042 | t-test | 0.098957 |
| *E-B-Farnesene* | 2V | 0.243583 | t-test | 0.000609 |
| *E-B-Farnesene* | -0.5V | 0.330473 | t-test | 0.662343 |
| *E-B-Farnesene* | 1V | 0.269318 | t-test | 0.231994 |
| *E-B-Farnesene* | -4V | 0.014306 | Wilcoxon test | 0.983548 |
| *E-B-Farnesene* | -1V | 0.015806 | Wilcoxon test | 0.606134 |
| *Nepetalactone* | 0V | 0.241889 | t-test | 0.442483 |
| *Nepetalactone* | 8V | 0.000263 | Wilcoxon test | 0.894499 |
| *Nepetalactone* | 0.5V | 0.092707 | t-test | 0.054763 |
| *Nepetalactone* | -8V | 0.134296 | t-test | 0.028586 |
| *Nepetalactone* | -4V | 0.028697 | Wilcoxon test | 0.140126 |
| *Nepetalactone* | -1V | 0.449722 | t-test | 0.590842 |
| *Nepetalactone* | 1V | 0.796161 | t-test | 0.610557 |
| *Nepetalactone* | 4V | 0.321075 | t-test | 0.48961 |
| *Nepetalactone* | -2V | 0.029429 | Wilcoxon test | 0.710103 |
| *Nepetalactone* | -0.5V | 0.334263 | t-test | 0.025636 |
| *Nepetalactone* | 2V | 0.320976 | t-test | 0.688976 |
| | *Aphid* | | | |
| *E-B-Farnesene* | 0V | 0.438858 | t-test | 0.442166 |
| *E-B-Farnesene* | -2V | 0.03087 | Wilcoxon test | 0.76058 |
| *E-B-Farnesene* | 8V | 4.82E-05 | Wilcoxon test | 0.000539 |
| *E-B-Farnesene* | 4V | 0.188464 | t-test | 0.012352 |
| *E-B-Farnesene* | 0.5V | 0.001081 | Wilcoxon test | 0.930609 |
| *E-B-Farnesene* | -8V | 0.086922 | t-test | 0.024167 |
| *E-B-Farnesene* | 2V | 0.336856 | t-test | 0.710789 |
| *E-B-Farnesene* | -0.5V | 0.491055 | t-test | 0.284925 |
| *E-B-Farnesene* | 1V | 0.008537 | Wilcoxon test | 0.527793 |

| | | | | |
|---|---|---|---|---|
| *E-B-Farnesene* | -4V | 0.226394 | t-test | 0.03314 |
| *E-B-Farnesene* | -1V | 0.535793 | t-test | 0.148855 |
| *Linalool* | 0V | 0.552438 | t-test | 0.422305 |
| *Linalool* | -2V | 0.835308 | t-test | 0.324065 |
| *Linalool* | 8V | 0.302913 | t-test | 0.018379 |
| *Linalool* | 4V | 0.076378 | t-test | 0.028693 |
| *Linalool* | 0.5V | 0.000302 | Wilcoxon test | 0.383957 |
| *Linalool* | -8V | 0.065474 | t-test | 0.024847 |
| *Linalool* | 2V | 0.785743 | t-test | 0.300126 |
| *Linalool* | -0.5V | 0.321191 | t-test | 0.873592 |
| *Linalool* | 1V | 0.24153 | t-test | 0.997715 |
| *Linalool* | -4V | 0.636963 | t-test | 0.260922 |
| *Linalool* | -1V | 0.139465 | t-test | 0.944801 |
| *E2Heptenal* | 4V | 0.400715 | t-test | 0.369161 |
| *E2Heptenal* | -4V | 0.065556 | t-test | 0.203079 |
| *E2Heptenal* | -1V | 0.974377 | t-test | 0.805237 |
| *E2Heptenal* | 2V | 0.011034 | Wilcoxon test | 0.070806 |
| *E2Heptenal* | 0.5V | 0.290664 | t-test | 0.8103 |
| *E2Heptenal* | -2V | 0.294889 | t-test | 0.157121 |
| *E2Heptenal* | -0.5V | 0.534845 | t-test | 0.779548 |
| *E2Heptenal* | 8V | 1.13E-05 | Wilcoxon test | 0.010218 |
| *E2Heptenal* | 0V | 0.445344 | t-test | 0.346146 |
| *E2Heptenal* | -8V | 0.011554 | Wilcoxon test | 0.001857 |
| *E2Heptenal* | 1V | 0.490286 | t-test | 0.023695 |
| | | *Fly* | | |
| *Linalool* | 0V | 0.112444 | t-test | 0.766105 |
| *Linalool* | -2V | 0.129577 | t-test | 0.238119 |
| *Linalool* | 8V | 0.390894 | t-test | 0.364268 |
| *Linalool* | 4V | 0.226619 | t-test | 0.477244 |
| *Linalool* | 0.5V | 0.000112 | Wilcoxon test | 0.360236 |
| *Linalool* | -8V | 0.969582 | t-test | 4.08E-05 |
| *Linalool* | 2V | 0.287072 | t-test | 0.698808 |
| *Linalool* | -0.5V | 0.078759 | t-test | 0.39984 |
| *Linalool* | 1V | 0.036876 | Wilcoxon test | 0.655793 |
| *Linalool* | -4V | 0.218778 | t-test | 0.118187 |
| *Linalool* | -1V | 0.498357 | t-test | 0.222504 |
| *E2Hexenal* | -0.5V | 0.002647 | Wilcoxon test | 0.818406 |
| *E2Hexenal* | -2V | 0.359048 | t-test | 0.207099 |
| *E2Hexenal* | 0.5V | 0.131089 | t-test | 0.898475 |
| *E2Hexenal* | 8V | 0.5625 | t-test | 0.067113 |

| | | | | |
|---|---|---|---|---|
| *E2Hexenal* | 2V | 0.027542 | Wilcoxon test | 0.153171 |
| *E2Hexenal* | -8V | 0.158473 | t-test | 0.008099 |
| *E2Hexenal* | -1V | 0.260978 | t-test | 0.171092 |
| *E2Hexenal* | 1V | 0.232613 | t-test | 0.512644 |
| *E2Hexenal* | -4V | 0.002354 | Wilcoxon test | 0.015435 |
| *E2Hexenal* | 0V | 0.041663 | Wilcoxon test | 0.202343 |
| *E2Hexenal* | 4V | 0.037295 | Wilcoxon test | 0.013398 |
| *Limonene* | 0V | 0.017858 | Wilcoxon test | 0.491224 |
| *Limonene* | -2V | 0.241765 | t-test | 0.264184 |
| *Limonene* | 8V | 0.323905 | t-test | 0.050508 |
| *Limonene* | 4V | 0.002174 | Wilcoxon test | 0.574827 |
| *Limonene* | 0.5V | 0.000384 | Wilcoxon test | 0.898565 |
| *Limonene* | -8V | 0.002931 | Wilcoxon test | 9.59E-05 |
| *Limonene* | 2V | 0.113739 | t-test | 0.466148 |
| *Limonene* | -0.5V | 0.002656 | Wilcoxon test | 0.429328 |
| *Limonene* | 1V | 0.336068 | t-test | 0.286309 |
| *Limonene* | -4V | 0.002214 | Wilcoxon test | 0.00864 |
| *Limonene* | -1V | 0.840099 | t-test | 0.316622 |
