## supplemental text - specimen preparation and fixation methods for bioimaging for "Electrostatic facilitation of odorant capture in insects"

### Fixation and embedding in LR White for light microscopy

#### Materials

1. **Buffer**: 0.1M phosphate buffer pH 7.2 (stock)
2. **Buffer**: 0.05M phosphate buffer pH 7.2
3. **Fixative**: 4% paraformaldehyde+2.5% glutaraldehyde in 0.05M phosphate buffer
4. **Ethanol**
5. **Glass vials**
6. **Paper/card to write labels in pencil and place inside vial**
7. **Plastic capsules for embedding**
8. **Oven**
9. **Tweezers**

#### Method

1. **Prepare fixative** in *fume hood* as follows:

For 100ml:

- Weigh 4g paraformaldehyde powder into a 50ml conical (Falcon) tube with a screw top and add 30 ml $dH_2O$.
- Heat to approximately 70ºC (screw cap on loose) in a beaker of water on a hotplate. Check temperature regularly with a thermometer – **do not let it boil**.
- Once at 70ºC and the powder starts to dissolve, carefully add 1-4 drops of 1M NaOH using a Pasteur pipette until the solution becomes clear.
- Place the tube on ice and allow to cool to room temperature.
- Add Glutaraldehyde (10ml of a 25% solution) and make up to 50ml with distilled water.
- Mix equal volumes of glut/paraformaldehyde mix and 0.1M phosphate buffer. Check pH 7.2 (might need slight adjustment with NaOH). Keep at 4ºC until use.

2. **Fix** - 2h at RT with rotation and then at 4ºC overnight. Samples can be placed under a mild vacuum initially to ensure fixative infiltration.
3. **Wash** - 3 x 0.05M phosphate buffer – 30min in each change. Samples can be stored in fridge in buffer for a few days.
4. **Dehydrate** in graded ethanol series (these times can be increased if needed depending on the sample type):

   - 10%EtOH rt 1x1hr
   - 20%EtOH rt 1x1hr
   - 30%ETOH rt 1x1hr

- 40%EtOH rt 1x1hr
- 50%EtOH rt 1x1hr
- 60%EtOH rt 1x1hr
- **70%EtOH 4°C overnight**
- 80% EtOH 2x1hr
- 90%EtOH 2x1hr
- 100% dry ETOH 3x1hr

5. **Infiltration** in increasing conc. of resin. The times can be increased, especially if dealing with large and/or hard samples. A mild vacuum can help infiltration when samples are in pure resin.

    - Ethanol 100% : LRWhite 4:1 1hr
    - Ethanol 100% : LRWhite 3:2 1hr
    - Ethanol 100% : LRWhite 2:3 1hr
    - Ethanol 100% : LRWhite 1:4 1hr
    - 100% LRWhite 1hr then 4°C overnight.
    - 100% LRWhite 1hr at RT with rotation.

6. **Embedding** – place samples in capsules, fill capsule with fresh resin, leave the lid open and place in oven. Run the nitrogen gas tap for about 15-30min to remove oxygen from oven. Polymerise @ 50 or 60 °C for 24h (50°C is better for immuno). Once polymerised, let samples cool down before sectioning. If resin is still a bit ‘sticky’ you can place the samples in the oven (run nitrogen gas for 15 min again) for another hour.
